# Helicase-deficient TFIIH causes severe disease features via persistent DNA excision without damage removal

**DOI:** 10.64898/2026.09.05.749590

**Authors:** David Häckes, Arjan F. Theil, Geun Hoe Kim, Youngmin Kim, Paula J. van der Meer, Céline Van Wassenhove, Qing Yu, Karen L. Thijssen, Anja Raams, Vera van Buren, Wim Vermeulen, Martijn S. Luijsterburg, Jurgen A. Marteijn, Jun-Hyuk Choi, Hannes Lans

**Author notes:** **Corresponding author:** Hannes Lans. These authors contributed equally to this work (David Häckes, Arjan F. Theil).

## Abstract

Nucleotide excision repair (NER) removes helix-distorting DNA lesions through the ten-subunit TFIIH complex, whose XPB and XPD translocase/helicase activities unwind DNA to enable damage verification and subsequent endonucleolytic DNA incisions. While most XPD mutations cause xeroderma pigmentosum, specific helicase-deficient mutations cause severe Cockayne syndrome (CS) features, including progressive neurodegeneration, for which the basis remains unclear. Here we show that loss of XPD helicase activity traps TFIIH in a futile repair cycle in which DNA is incised at the wrong position, leading to repeated DNA excision and resynthesis without removal of the lesion. Using *C. elegans*, we find that this futile DNA excision cycle produces severe neuronal dysfunction *in vivo* that depends on transcription-coupled NER activity and is rescued by preventing recruitment of helicase-deficient TFIIH. These findings demonstrate that NER incisions can occur without XPD-mediated damage verification and that persistent futile DNA excision cycles cause severe disease features, indicating that persistent NER intermediates are more pathogenic than unrepaired DNA lesions.

## 2 Introduction

Maintaining genomic stability requires cells to contend with a continuous stream of DNA lesions generated by both endogenous and environmental sources. Left unrepaired, these lesions can drive mutagenesis, leading to cancer, and premature aging, which is why cells rely on an intricate network of DNA repair and DNA damage signaling pathways to detect and remove diverse types of lesions^1^. Among these pathways, nucleotide excision repair (NER) is responsible for removing a broad spectrum of bulky and helix-distorting DNA lesions, including those induced by UV radiation and various chemical mutagens^2,3^. Central to this pathway is the ten-subunit transcription factor IIH (TFIIH) complex, which acts in both transcription initiation and NER^4–6^. In NER, TFIIH is recruited to damaged DNA following lesion detection by either of two independent mechanisms. In global genome NER (GG-NER), DNA damage is recognized by the XPC-RAD23B-CETN2 complex, which constantly probes the entire genome for structural distortions and recruits TFIIH upon stable DNA damage binding^7–9^. In transcription-coupled NER (TC-NER), DNA damage in the transcribed strand of active genes is recognized by the stalling of the elongating RNA Polymerase II (Pol II) complex at a lesion^10^. This stalling triggers the sequential recruitment of the translocase CSB and the E3 ubiquitin ligase complex CRL4*^CSA^*, which ubiquitylates Pol II to facilitate both its removal from the lesion and the recruitment of downstream repair factors. This is followed by binding of the UVSSA-USP7 complex, which, together with STK19, subsequently recruits and positions TFIIH^11–14^. TFIIH then opens the DNA around the lesion through the translocase activity of its XPB subunit and the translocase/helicase activity of its XPD subunit^15^. In particular, XPD translocates along the damaged single-stranded DNA in the 5*^′^ →* 3*^′^* direction until its progression is blocked by the lesion^16–20^. This activity is stimulated by XPG and by the scaffold factor XPA, while RPA coats the undamaged strand, together stabilizing the incision complex and promoting efficient lesion verification^17,18,21^. The arrest of XPD at the lesion serves as the final verification of the damage and licenses the subsequent DNA incisions 5*^′^* and 3*^′^* of the lesion, respectively, by the ERCC1-XPF and XPG endonucleases to excise a damaged ssDNA segment of 25–30 nt^22–24^. Following the removal of this damaged ssDNA segment together with TFIIH, the resulting ssDNA gap is filled in by novel DNA synthesis by DNA polymerases and sealed by ligation^25,26^.

Hereditary defects in NER give rise to a group of rare autosomal recessive disorders. Defects in GG-NER cause xeroderma pigmentosum (XP), which is primarily characterized by extreme photosensitivity and a strong predisposition to UV-induced skin cancer, and, particularly in case of XPA mutations, progressive neurodegeneration due to accumulation of mutagenic lesions^27^. Defects in the TC-NER-specific factors CSB and CSA mostly lead to Cockayne syndrome (CS), defined by photosensitivity, severe growth failure, segmental progeria and progressive neurodegeneration^28^, while mutations in UVSSA cause the clinically milder UV-sensitive syndrome (UV^S^S), which also presents with photosensitivity but lacks the severe growth and neurodevelopmental features of CS^29–31^. Intriguingly, different mutations in TFIIH subunits, as well as in ERCC1, XPF and XPG, can give rise to either XP or the combined XP/Cockayne syndrome complex (XPCS; in its most severe form also referred to as cerebro-oculo-facio-skeletal syndrome)^4,32^. Additionally, some mutations in TFIIH subunits cause photosensitive trichothiodystrophy (TTD; also referred to as XP/TTD). TTD is largely attributed to transcriptional insufficiency caused by TFIIH instability^33,34^, as TFIIH also serves as a general transcription factor required for transcription initiation by Pol II. The molecular pathogenesis of CS has for a long time been less well understood. However, we and others have shown that defects in CSB or CSA lead to a defect in the repair of transcription-blocking DNA-protein crosslinks and the inability to remove stalled Pol II from transcription-blocking DNA lesions^31,35–40^. Furthermore, we previously showed that severe defects in XPF or XPG cause TFIIH to become persistently stalled in the absence of DNA incision, forming toxic repair intermediates^21,41^. Together, these findings suggest that persistently stalled NER intermediates trigger a global transcription shutdown, which drives the cellular dysfunction and progressive neurodegeneration observed in CS^14^.

Thus far, it has remained unclear why some mutations in TFIIH subunits, particularly in *XPD/ERCC2*, give rise to CS features in addition to XP, and whether this similarly involves the persistence of a repair intermediate. To address this, we compared the molecular NER mechanism in isogenic cells carrying different XPD patient mutations associated with different diseases: G47R, which causes XPCS^42,43^, S541R, which causes XP^44–46^, and L485P, which causes XP/TTD^47^. Here, we demonstrate that the inability of XPD to translocate along DNA, caused by disruption of its helicase activity due to the G47R mutation, leads to a persistent futile DNA repair cycle in which both NER incisions are made on the same side, rather than on both flanks, of the actual lesion. Using *C. elegans* as a model system^48^, we demonstrate *in vivo* that these persistent, unproductive DNA repair cycles further exacerbate the already severe neuronal dysfunction caused by impaired repair. Strikingly, this exacerbated neuronal dysfunction is attenuated when the recruitment of mutant TFIIH is prevented.

## 3 Results

### XPD patient mutations differentially impact cellular survival

To better understand why certain *XPD/ERCC2* mutations cause CS features, we generated isogenic cell lines carrying different XPCS- or XP-associated patient mutations and compared their molecular NER mechanism and cellular phenotype. We mapped these mutations onto a cryo-EM structure of XPD and TFIIH^18^, in which the evolutionary conservation of XPD residues is color coded (see Methods, Figure 1a). We selected the XPCS-causing G47R mutation^42,43^, which is located in the conserved ATP-binding domain of XPD and results in a helicase-dead protein^49,50^. We compared this with the XP-causing S541R mutation^44–46^, located at the conserved ssDNA entry pore, and the XP/TTD-causing L485P mutation^47^, which is located in a less conserved region near the XPD-XPB protein interface. To investigate their impact on NER activity, and in particular on TFIIH function, all three point mutations were introduced by CRISPR-Cas9 into the *XPD/ERCC2* gene of previously generated GFP-XPB and CSB-mClover knock-in (KI) U2OS cells^21,51^. Clonal cell lines were established and homozygous mutations confirmed by sequencing, for both GFP-XPB and CSB-mClover cell lines (Figure S1a–b). Confocal microscopy and quantification of GFP fluorescence in the GFP-XPB cell lines confirmed that TFIIH was expressed at comparable levels in all mutant and parental cell lines (Figure S1c, Figure 1b).

**Figure 1.**
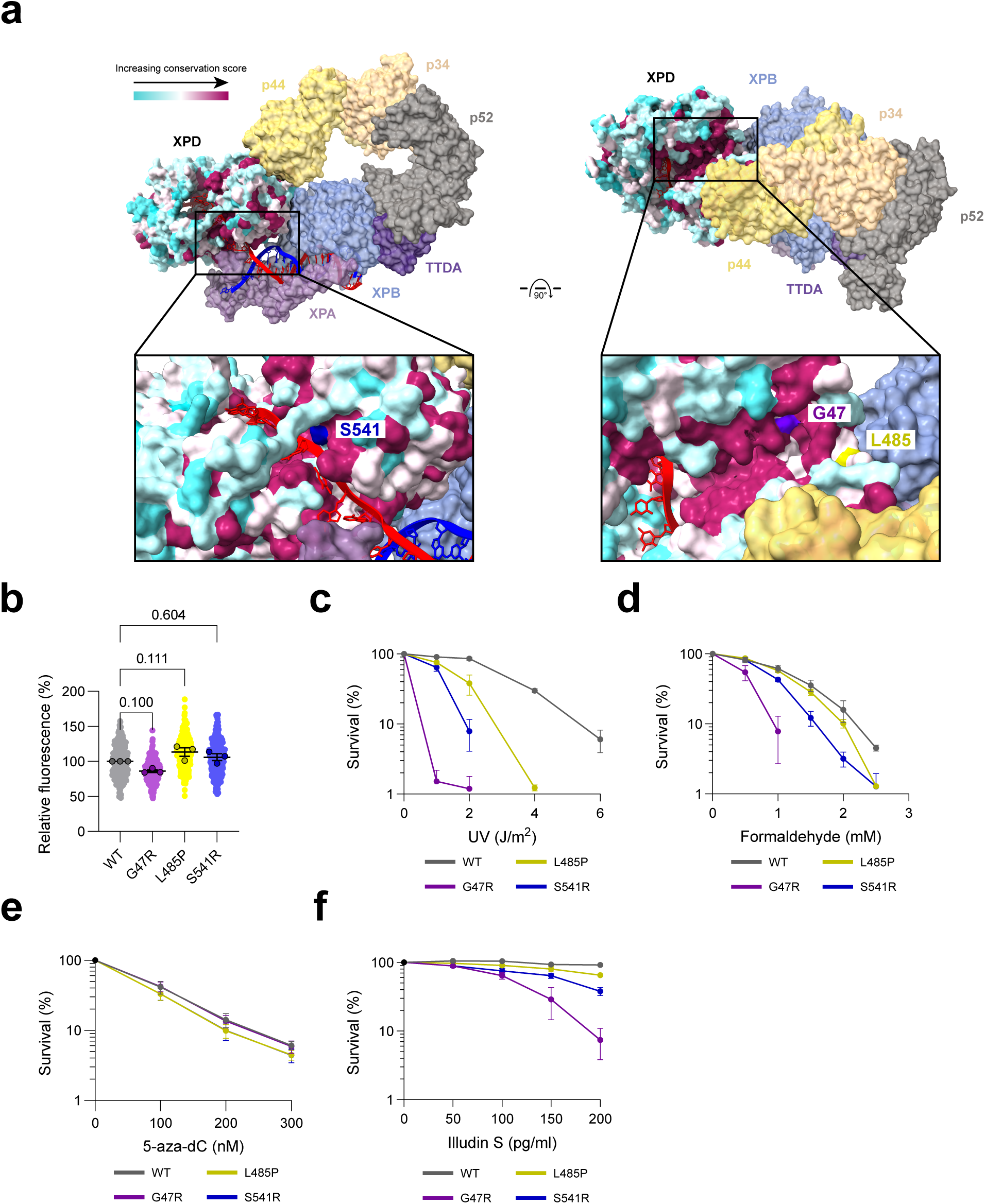
XPD patient mutations differentially impact cellular survival. **A** Surface representation of TFIIH (PDB: 6RO4) with the XPD subunit colored according to evolutionary conservation across *H. sapiens*, *M. musculus*, *B. taurus*, *A. thaliana*, *S. cerevisiae* and *C. elegans* XPD orthologs, from variable (cyan) to highly conserved (magenta). The positions of the patient mutations G47R, L485P and S541R are indicated. **B** Mean nuclear GFP-XPB fluorescence intensity in parental (WT) and XPD-G47R, -L485P and - S541R GFP-XPB knock-in U2OS cells, normalized to WT. n = 285 (WT), 279 (G47R), 252 (L485P) and 265 (S541R) cells from three independent experiments. Numbers represent p-values (one-way ANOVA corrected for multiple comparisons). **C, D** Clonogenic survival of parental (WT) and XPD-G47R, -L485P and -S541R CSB-mClover knock-in U2OS cells after UV-C (**C**) or formaldehyde (FA, **D**) treatment. **E, F** Clonogenic survival of the same cell lines after 5-aza-2*^′^*-deoxycytidine (5-Aza-dC, **E**) or illudin S (**F**) treatment. Mean and s.e.m. of three (**C–E**) or two (**F**) independent experiments, each performed in technical triplicate. Source data are provided as a Source Data file.

We performed clonogenic survival assays to functionally test how each of the XPD mutations affected cellular survival after DNA damage induction. Following exposure to UV-C irradiation, we observed that XPD-G47R mutant CSB-mClover and GFP-XPB cells were the most hypersensitive (Figure 1c, Figure S1d). XPD-L485P and XPD-S541R mutant cells both displayed an intermediate hypersensitivity compared to wild type cells, with XPD-L485P mutant cells being the least hypersensitive. A similar pattern was observed after treatment with the cross-linking agent formaldehyde (FA), which induces both DNA crosslinks as well as DNA-protein crosslinks (DPCs)^52^. XPD-G47R mutant cells were the most hypersensitive to FA exposure, while XPD-S541R mutant cells displayed an intermediate hypersensitivity, and XPD-L485P mutant cells were as sensitive as the wild type control cells (Figure 1d). To determine whether this FA hypersensitivity was due to DNA-DNA or DNA-protein crosslinks, we tested survival after exposure to 5-aza-2*^′^*-deoxycytidine (5-Aza-dC), which exclusively creates DPCs by covalently trapping DNA methyltransferase DNMT1 to DNA^53^. All mutant cell lines exhibited comparable sensitivity to 5-Aza-dC as wild type cells (Figure 1e), indicating that the differential response to FA is primarily driven by the processing of DNA crosslinks rather than DPCs. Finally, we tested the sensitivity to illudin S, which generates DNA damage specifically processed by TC-NER^54^. This again showed that XPD-G47R mutant cells are the most hypersensitive to DNA damage processed by NER (Figure 1f). Together, these results indicate that the XPD mutation associated with the most severe disease phenotypes, i.e. G47R, also leads to the most severe functional defects in cells after DNA damage induction.

### XPD-G47R does not cause persistent TFIIH or Pol II stalling

Previously, we showed that persistent binding of TFIIH to DNA damage, as observed in XPCS cells deficient in either ERCC1-XPF or XPG, contributes to severe cellular dysfunction as characteristic for CS^21,41^. To investigate whether TFIIH similarly stalls at UV-induced DNA damage in XPD-G47R mutant cells, we tested Fluorescence Recovery After Photobleaching (FRAP) in the mutant GFP-XPB U2OS cells. In this FRAP experiment, the incomplete fluorescence recovery in a photobleached subnuclear area reflects the immobilization, i.e. binding, of GFP-tagged TFIIH molecules to DNA damage during NER^55–57^. We observed a clear immobilization of GFP-XPB/TFIIH in wild type parental cells immediately after UV irradiation, which was absent upon knockout of XPC, confirming dependence on GG-NER (Figure 2a, Figure S2a). This was also observed in XPD-L485P mutant cells. However, the XPD-S541R cells and, in particular, the XPD-G47R mutant cells displayed a significantly reduced immobile fraction of TFIIH compared to the parental cells (Figure 2a, Figure S2b–d), indicating that these mutant TFIIH complexes associate less efficiently or less stably with DNA damage.

**Figure 2.**
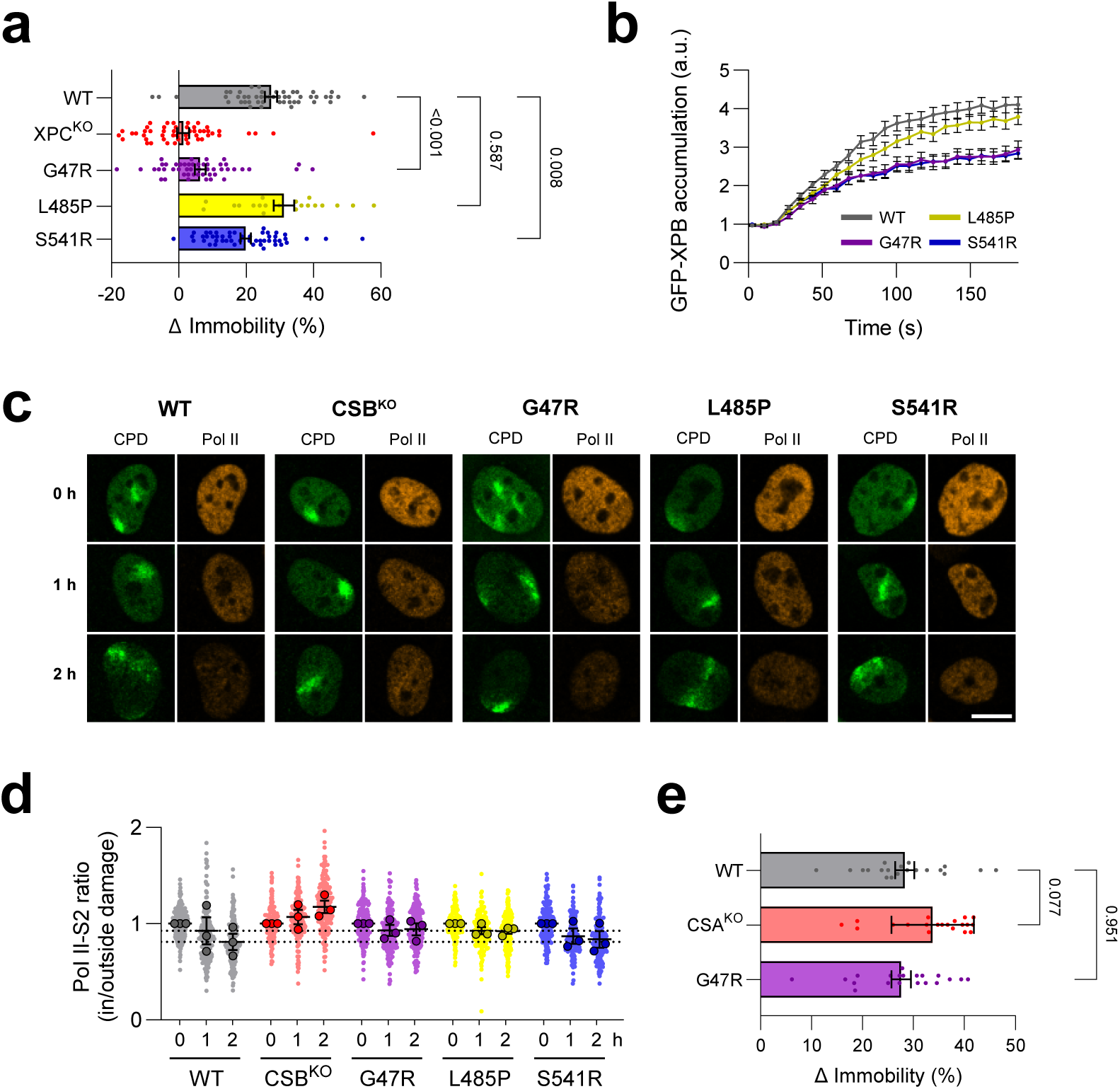
XPD-G47R does not cause persistent TFIIH or Pol II stalling. **A** Quantification of the UV-induced change in GFP-XPB immobile fraction as measured by FRAP in parental (WT), XPC knockout, and XPD-G47R, -L485P and -S541R GFP-XPB knock-in U2OS cells before and 5–30 min after 10 J/m^2^ UV-C irradiation. FRAP curves are shown in Figure S2a–d. Data pooled from five independent experiments, except L485P from two independent experiments; n = 50 (WT), 50 (XPC knockout), 49 (G47R), 20 (L485P) and 50 (S541R) cells. **B** Average accumulation of GFP-XPB at sites of local UV damage in parental (WT) and XPD-G47R, -L485P and -S541R GFP-XPB knock-in U2OS cells, induced using a 266 nm UV-C laser. Fluorescence intensities are background-corrected and normalized to pre-damage levels. n = 37 (WT), 40 (G47R), 38 (L485P) and 36 (S541R) cells from three independent experiments. **C** Representative immunofluorescence images of the conditions in (**D**), stained for CPD and Pol II-S2P. Scale bar, 10 *µ*m. **D** Quantification of elongating Pol II clearance from sites of local UV damage in parental (WT), XPD-G47R, -L485P, -S541R and CSB knockout CSB-mClover knock-in U2OS cells, untreated or treated with 100 *µ*M DRB for 1 or 2 h after local UV-C irradiation (100 J/m^2^) through a 5 *µ*m pore filter. Plotted is the mean ratio of Pol II-S2P signal inside versus outside the locally damaged area, defined by CPD staining. n (0, 1 and 2 h) = 203, 193 and 215 (WT); 227, 197 and 208 (G47R); 187, 185 and 215 (L485P); 275, 163 and 185 (S541R); and 215, 187 and 190 (CSB knockout) cells, from three independent experiments. **E** Quantification of the UV-induced change in CSB-mClover immobile fraction as measured by FRAP in parental (WT), XPD-G47R and CSA knockout CSB-mClover knock-in U2OS cells before and 5–30 min after 10 J/m^2^ UV-C irradiation. FRAP curves are shown in Figure S2e. Data pooled from two independent experiments; n = 20 UV-irradiated cells per cell line, each compared to the mean of the corresponding untreated condition (n = 20 (WT), 23 (XPD-G47R) and 19 (CSA knockout) cells). Mean and s.e.m. are shown. Numbers represent p-values (one-way ANOVA corrected for multiple comparisons). Source data are provided as a Source Data file.

To investigate whether the initial recruitment of mutant TFIIH to DNA damage was compromised, we monitored real-time GFP-XPB DNA damage accumulation using live cell imaging after DNA damage induction with a 266 nm UV-C laser. We observed an immediate accumulation of wild type TFIIH and an almost similar accumulation of XPD-L485P mutant XPD (Figure 2b). By comparison, XPD-S541R and XPD-G47R mutant TFIIH also evidently accumulated, but showed a lower peak accumulation. Together, these data indicate that the XPD-G47R mutation does not lead to persistent TFIIH stalling, but rather causes TFIIH to be recruited and bound less efficiently to DNA damage.

In addition to TFIIH, the persistent stalling of Pol II at DNA damage, due to its impaired clearance, has been implicated in CS pathogenesis due to CSB and CSA deficiency^31,35–37^. Therefore, we assessed the clearance of Pol II from DNA damage by immunofluorescence of elongating Pol II at sites of local UV damage after transcription inhibition^58^. As previously noted^59^, we observed an accelerated loss of the elongating Pol II signal at sites of local UV damage in wild type cells, but not in CSB knockout U2OS cells, which were included as control (Figure 2c–d). This indicates that Pol II is efficiently cleared from DNA damage in wild type cells, but remains stalled in the absence of CSB activity. We observed a similar loss of Pol II signal in XPD-S541R mutant cells as in wild type cells, but not in XPD-L485P and XPD-G47R mutant cells, although their defect was less severe than the complete absence of clearance observed in CSB knockout cells (Figure 2d). As the XPD-G47R mutation was shown to inactivate the XPD helicase activity *in vitro*^49,50^, this indicates that XPD helicase activity promotes Pol II clearance. This was recently also concluded using the same assay and K48R and G675R helicase-dead mutant XPD proteins, while it was also shown that activity of the ubiquitin-dependent segregase VCP serves as a backup mechanism to remove Pol II from DNA damage^59^. We next measured FRAP of CSB-mClover after DNA damage induction as complementary approach to assess Pol II stalling, as the immobilization of CSB in FRAP is a direct measure of the binding of elongating Pol II to DNA damage^37,51^. This showed that following UV irradiation, XPD-G47R mutant cells exhibit a CSB immobile fraction comparable to that of wild type cells, whereas CSA knockout cells, which were included as control, showed a clearly increased CSB immobilization (Figure 2e, Figure S2e). Together, these results suggest that Pol II is still cleared in XPD-G47R mutant cells and is not persistently stalled at DNA damage sites as observed in CSA- or CSB-deficient cells. This indicates that neither persistent TFIIH stalling nor persistent Pol II stalling explains the severe phenotype associated with the XPD-G47R mutation, suggesting the involvement of a different pathological mechanism.

### Helicase-dead XPD initiates DNA repair synthesis without removal of the lesion

We next examined how the different XPD mutations affected the NER pathway itself. Using immunofluorescence in the XPD mutant CSB-mClover KI cells, we found that the NER incision complex factors XPA, XPF, XPG and RPA32 were all visibly recruited to sites of local UV damage in each mutant cell line, indicating that the incision complex still assembles in the presence of these XPD mutants (Figure 3a). Therefore, we measured DNA synthesis during NER, using the Unscheduled DNA Synthesis (UDS) assay^60^ by quantifying the incorporation of 5-ethynyl-2*^′^*-deoxyuridine (EdU) after global UV irradiation in wild type and XPD mutant CSB-mClover KI U2OS cells. This showed that the parental cells exhibited a robust UDS signal, indicative of efficient NER (Figure 3b–c). XPD-L485P and XPD-S541R mutant cells showed a significantly reduced, but still clearly detectable residual UDS signal, in line with their intermediate UV hypersensitivity (Figure 1c, Figure S1d). Strikingly, however, XPD-G47R mutant cells, which were extremely UV hypersensitive (Figure 1c, Figure S1d), also exhibited a clearly detectable residual UDS signal, to the same level as XPD-S541R mutant cells. Independently, this was also observed in the XPD-G47R mutant GFP-XPB KI U2OS cells (Figure S3a). To determine whether this UV-induced UDS reflects NER activity, we depleted the essential NER endonuclease XPF using siRNA, and found that this completely abrogated the UDS signal in both wild type and XPD-G47R mutant cells (Figure 3d).

**Figure 3.**
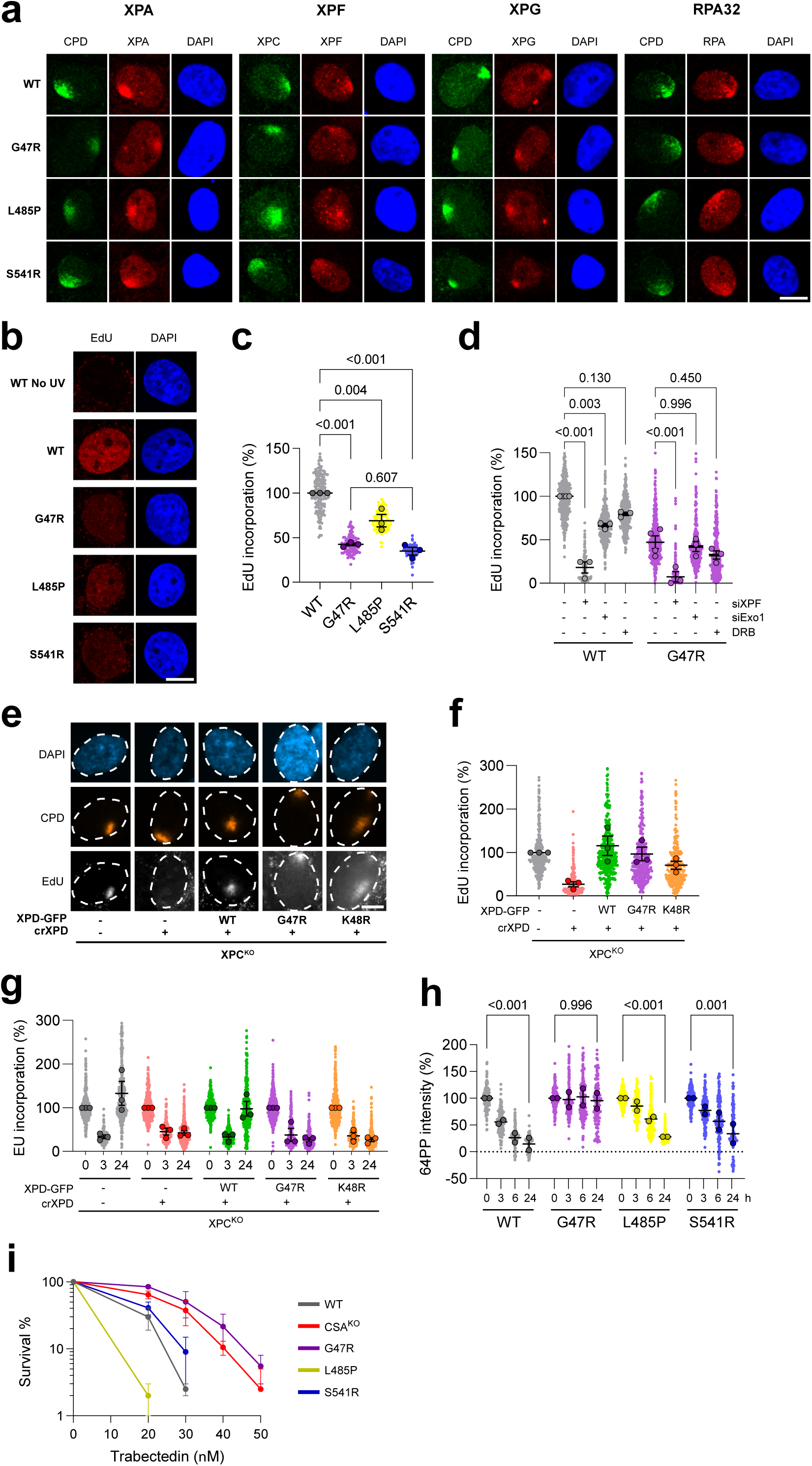
Helicase-dead XPD initiates DNA repair synthesis without removal of the lesion. **A** Representative immunofluorescence images of XPA, XPF, XPG and RPA32 at sites of local UV-C damage (100 J/m^2^ through a 5 *µ*m pore filter) in parental (WT) and XPD-G47R, -L485P and -S541R CSB-mClover knock-in U2OS cells, with locally damaged regions identified by CPD co-staining, except for the XPF panels, in which XPC co-staining was used to identify the damaged regions. **B** Representative immunofluorescence images of the unscheduled DNA synthesis (UDS) quantified in (**C**), showing EdU incorporation and DAPI. **C** UDS measured by EdU incorporation for 1 h after 8 J/m^2^ UV-C irradiation in parental (WT) and XPD-G47R, -L485P and -S541R CSB-mClover knock-in U2OS cells. EdU intensities are normalized to WT. n = 390 (WT), 222 (G47R), 215 (L485P) and 230 (S541R) cells from three independent experiments. **D** UDS as in (**C**) in parental (WT) and XPD-G47R cells transfected with siCtrl, siXPF or siExo1, or treated with 100 *µ*M DRB. n = 474 (WT, siCtrl), 307 (WT, siXPF), 422 (WT, siExo1), 392 (WT, DRB), 533 (G47R, siCtrl), 331 (G47R, siXPF), 380 (G47R, siExo1) and 376 (G47R, DRB) cells, from four (siCtrl) or three (siXPF, siExo1, DRB) independent experiments. **E** Representative images of the TC-NER-dependent UDS quantified in (**F**), displaying EdU incorporation at sites of local UV-C damage identified by CPD co-staining. **F** TC-NER-dependent UDS, measured by EdU incorporation for 4 h immediately after local UV-C irradiation through a 5 *µ*m pore filter (100 J/m^2^) in XPC knockout (XPC^KO^) hTERT-RPE1 cells stably expressing inducible Cas9; endogenous XPD was silenced by transfection of a synthetic crRNA (crXPD) and cells were complemented with wild type (WT), G47R or K48R XPD-GFP as indicated. Locally damaged regions were identified by CPD co-staining. EdU intensities are normalized to WT XPD-GFP. n = 386 (XPC^KO^), 255 (crXPD), 354 (crXPD + WT XPD-GFP), 378 (crXPD + G47R) and 288 (crXPD + K48R) cells, from three independent experiments. **G** Recovery of RNA synthesis (RRS), measured by 5-ethynyluridine (EU) incorporation at 0, 3 and 24 h after 12 J/m^2^ UV-C irradiation, in cells as in (**F**). EU intensities are normalized to the untreated control of each cell line. n (0, 3 and 24 h) = 702, 677 and 442 (XPC^KO^); 455, 455 and 410 (crXPD); 753, 591 and 404 (crXPD + WT XPD-GFP); 791, 647 and 410 (crXPD + G47R); and 713, 505 and 467 (crXPD + K48R) cells, from three independent experiments. **H** 6-4PPs immunofluorescence signal at 0, 3, 6 and 24 h after 4 J/m^2^ UV-C irradiation in parental (WT) and XPD-G47R, -L485P and -S541R CSB-mClover knock-in U2OS cells, normalized to the signal at 0 h. n (0, 3, 6 and 24 h) = 191, 169, 156 and 140 (WT); 214, 190, 192 and 139 (G47R); 257, 189, 181 and 122 (L485P); and 215, 206, 200 and 128 (S541R) cells, from two independent experiments. **I** Clonogenic survival after trabectedin treatment of parental (WT), XPD-G47R, -L485P and -S541R, and CSA knockout CSB-mClover knockin U2OS cells, from two independent experiments, each in technical triplicate. Scale bars, 10 *µ*m. Mean and s.e.m. are shown. Numbers represent p-values (one-way ANOVA corrected for multiple comparisons). Source data are provided as a Source Data file.

These results are in line with previous findings showing residual UDS levels in extremely UV hypersensitive XPCS patient fibroblasts that harbor helicase-dead G675R or G602D XPD mutations^61^ and in XPD fibroblasts with overexpression of the helicase-dead K48R XPD mutant^62^. Indeed, we also found residual UDS in these and another (XPCS1PV, carrying R666W) XPCS patient fibroblasts expressing helicase-dead XPD^63,64^ (Figure S3b). Previously, fibroblasts from XPCS patients with helicase-dead XPD mutations, i.e. G47R, R666W, G675R, and G602D, were shown to accumulate DNA breaks after UV, which was reported to depend on transcription^63,65^. However, we did not find that the UDS signal in our XPD-G47R mutant CSB-mClover U2OS cells was largely dependent on transcription (Figure 3d). Also, it was suggested that residual repair synthesis in XPCS fibroblasts could be due to strand displacement synthesis, involving processing of the incised DNA by the exonuclease activity of EXO1^66^. We recently already showed that UV-induced DNA repair synthesis in wild type cells is partially dependent on EXO1^67^. However, we did not observe that the residual UDS in XPD-G47R mutant cells was any more dependent on EXO1 than in wild type cells (Figure 3d). Together, these results show that the XPD mutant cells retain a significant, NER-dependent capacity for repair synthesis.

UV-induced UDS mainly reflects GG-NER activity and not TC-NER activity^68^, as is also evident from the minimal effect of transcription inhibition (Figure 3d). As CS symptoms are specifically associated with TC-NER defects, we therefore also measured TC-NER-induced UDS in an alternative cellular model system. We used previously generated hTERT-RPE1 cells stably expressing inducible Cas9 in which XPC was knocked out to abolish GG-NER activity^69^. In these cells, wild type or mutant XPD-GFP was stably transfected, while endogenous XPD was silenced by transfection of a synthetic guide RNA (crRNA), as described previously^59^. Expression of the XPD-GFP variants and depletion of endogenous XPD were confirmed by immunoblotting (Figure S3c). In addition to G47R, we included the K48R helicase-dead XPD mutant^62,70^. We then measured UDS levels at sites of local UV damage and found that cells expressing wild type XPD or either XPD mutant exhibited a robust TC-NER-dependent UDS signal, whereas cells without XPD did not (Figure 3e–f). Next, we measured recovery of RNA synthesis (RRS) in these cells, which reflects the successful completion of TC-NER, by visualizing nascent transcription using 5-ethynyluridine incubation. Strikingly, this showed that while cells with wild type XPD normally recovered transcription after 24 h, cells without XPD or with helicase-dead XPD did not (Figure 3g). These results demonstrate that helicase-dead XPD mutant cells perform DNA repair synthesis, but that they still lack DNA repair capacity.

To directly measure DNA repair capacity by GG-NER in the XPD-L485P, XPD-S541R and XPD-G47R mutant cells, we monitored the removal of 6-4 photoproducts (6-4PPs) over time after UV irradiation, using immunofluorescence in CSB-mClover KI U2OS cells. As expected, 6-4PPs were efficiently removed within 6 h in the parental cells^71^. 6-4PPs were also removed in XPD-L485P and XPD-S541R mutant cells, albeit with slightly delayed kinetics, consistent with their reduced UDS signal. In stark contrast, 6-4PPs were not at all removed in XPD-G47R mutant cells, and DNA damage levels remained unchanged even at 24 h after UV irradiation (Figure 3h), indicative of severely impaired NER.

We next assessed TC-NER capacity by testing the clonogenic survival of the XPD mutant CSB-mClover KI U2OS cells following exposure to trabectedin. Trabectedin creates DNA monoadducts specifically detected and processed by TC-NER, but the trabectedin-DNA adducts block the NER endonuclease XPG at the incision step, creating toxic persisting DNA breaks generated by ERCC1-XPF. Consequently, TC-NER-proficient cells are hypersensitive to trabectedin, while TC-NER-deficient cells are resistant^72,73^. We found that the parental cells, as well as the XPD-L485P and XPD-S541R mutant cells were all hypersensitive to trabectedin, consistent with their capacity for DNA repair. In contrast, previously generated TC-NER-deficient CSA knockout U2OS cells^51^, as well as the XPD-G47R mutant cells, were highly resistant to trabectedin (Figure 3i). These results indicate that the XPD-G47R mutant cells do not process the trabectedin-DNA adducts normally and are therefore TC-NER-deficient. Together, our results indicate that XPD-G47R mutant cells exhibit significant UV-induced, GG- and TC-NER-dependent DNA repair synthesis, yet are completely DNA repair deficient. This suggests that in these cells a stretch of ssDNA might be excised and resynthesized that does not encompass the original DNA lesion.

### Helicase-dead XPD leads to misincision and continuous futile DNA excisions without damage removal

To directly test whether DNA excision occurs without the removal of DNA damage in XPD-G47R mutant cells, we purified the excised DNA fragments from UV-irradiated wild type parental and XPD-G47R mutant CSB-mClover KI U2OS cells and, additionally, determined whether these contained DNA damage using 6-4PP immunoprecipitation^71,74^. Purification of the fragments without 6-4PP immunoprecipitation revealed that, following UV irradiation, excised oligonucleotide fragments of approximately the same size were produced in the XPD-G47R mutant cells as in the parental cells, albeit at a reduced level (Figure 4a–b). No DNA fragments could be purified upon knockout of XPA in XPD-G47R mutant cells, showing that these DNA fragments are produced by NER activity. Next, we immunoprecipitated the excised DNA fragments with 6-4PP antibodies, which showed that the excised fragments in the wild type parental cells contained 6-4PPs, whereas the fragments in XPD-G47R mutant cells did not (Figure 4c–d). These data confirm that in XPD-G47R mutant cells NER is initiated and excises a normal-sized DNA fragment, but that it fails to remove any DNA damage.

**Figure 4.**
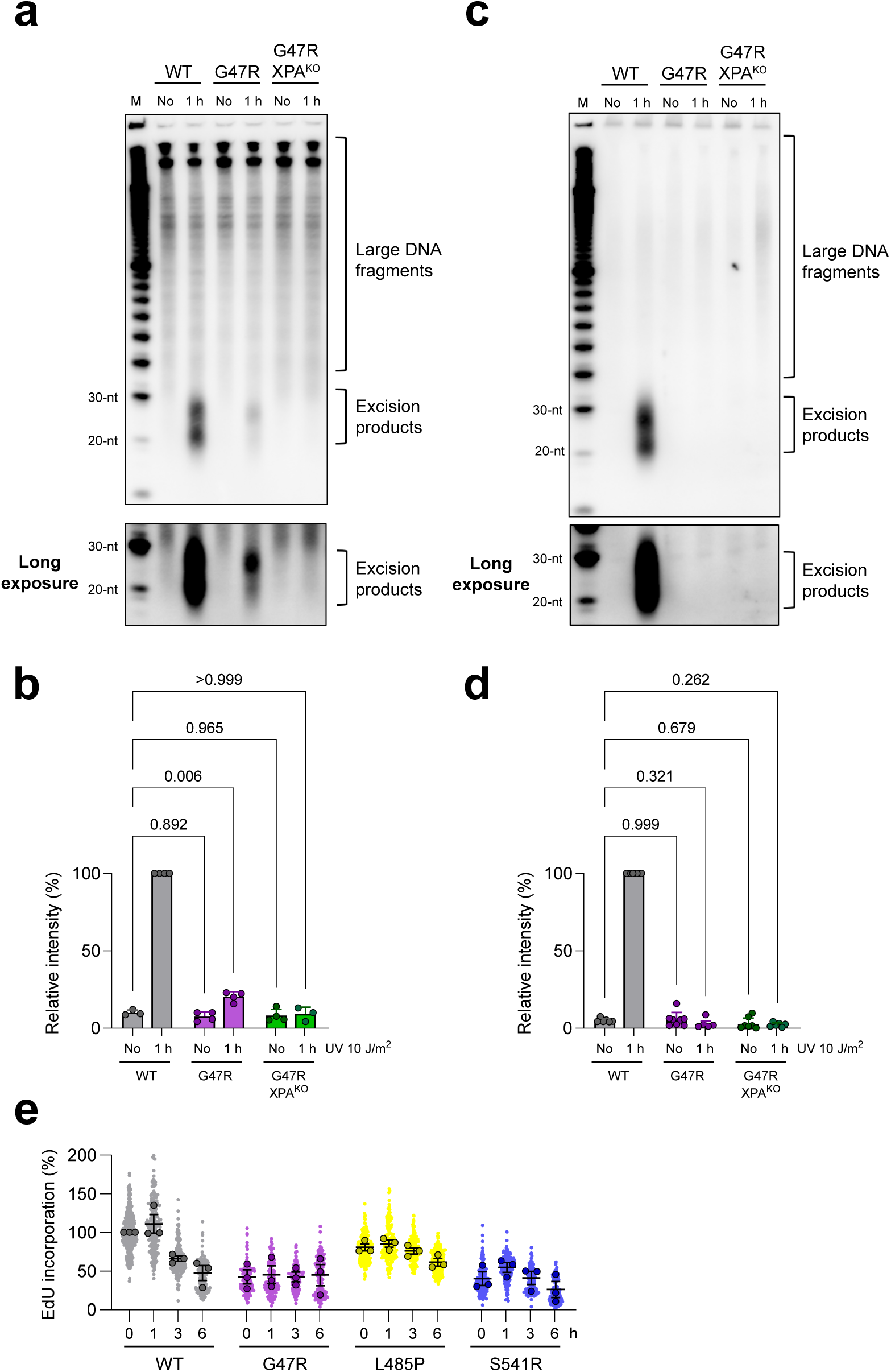
G47R mutant cells carry out futile DNA excision without damage removal. **A** *In vivo* excision assay in parental (WT), XPD-G47R and XPD-G47R/XPA KO double mutant CSB-mClover knock-in U2OS cells, before and 1 h after 10 J/m^2^ UV-C irradiation. Excised oligonucleotides were 3*^′^*-end labeled with iotin-11-dUTP, resolved by denaturing PAGE, and detected by streptavidin-HRP chemiluminescence. A representative experiment is shown. M denotes the size marker. **B** Quantification of signal intensities from (**A**), background-subtracted and normalized to the UV-irradiated WT. n (left to right) = 3, 4, 4, 4, 4 and 3 independently quantified membranes. **C** Immunoprecipitation of excised oligonucleotides from cells as in (**A**) with an anti-6-4PP antibody, detected as in (**A**). A representative experiment is shown. **D** Quantification of signal intensities from (**C**), background-subtracted and normalized to the UV-irradiated WT. n (left to right) = 6, 8, 8, 8, 8 and 8 independently quantified membranes. **E** UDS time course measured by EdU incorporation at 0, 1, 3 and 6 h after 8 J/m^2^ UV-C irradiation in parental (WT) and XPD-G47R, -L485P and -S541R CSB-mClover knock-in U2OS cells. EdU intensities were normalized to WT. n (0, 1, 3 and 6 h) = 406, 204, 222 and 207 (WT); 206, 207, 188 and 179 (G47R); 218, 190, 267 and 221 (L485P); and 216, 205, 199 and 181 (S541R) cells, from three independent experiments. Mean and s.e.m. are shown. Numbers represent p-values (one-way ANOVA corrected for multiple comparisons). Source data are provided as a Source Data file.

We reasoned that absence of damage removal should lead to a continuous futile cycle of DNA excision and resynthesis in XPD-G47R mutant cells. To investigate this, we measured UDS, which mainly reflects the repair of 6-4PPs^68^, at sequential time points up to 6 h after UV irradiation in parental and mutant CSB-mClover U2OS cells. In wild type, XPD-L485P, and XPD-S541R mutant cells the UDS levels peaked at 1 h after UV and then declined as repair was completed. In contrast, UDS in XPD-G47R mutant cells remained at a constant, sustained level throughout the entire course of time (Figure 4e), in line with a continuous cycle of futile DNA excision.

### Futile DNA excision mediated by helicase-dead XPD causes severe neuronal dysfunction *in vivo in C. elegans*

To determine whether this continuous futile DNA repair cycle is toxic to cells, in particular to differentiated post-mitotic cells *in vivo*, and thus can partly explain the more severe symptoms observed in patients with the XPD-G47R mutation, we utilized *C. elegans* as a multicellular *in vivo* model system. We and others have previously shown that *C. elegans* is excellently suited to investigate *in vivo* NER mechanisms and associated disease phenotypes^21,36,75–77^. Given the high degree of structural and sequence conservation between human XPD and worm XPD-1 (Figure 5a, Figure S4a), we generated mutant *C. elegans* strains with the corresponding amino acid substitutions using CRISPR-Cas9. Interestingly, while we successfully generated viable strains carrying G47R or L479P (corresponding to human L485P) mutations in *C. elegans xpd-1* (Figure 5b), we were unable to generate viable strains carrying the corresponding S541R mutation in *xpd-1*. A possible explanation is that the ssDNA entry pore appears more constricted in the *C. elegans* XPD-1 structural model than in human XPD, such that introducing an arginine at this position may fully occlude the pore in the worm and abolish an essential XPD-1 function.

**Figure 5.**
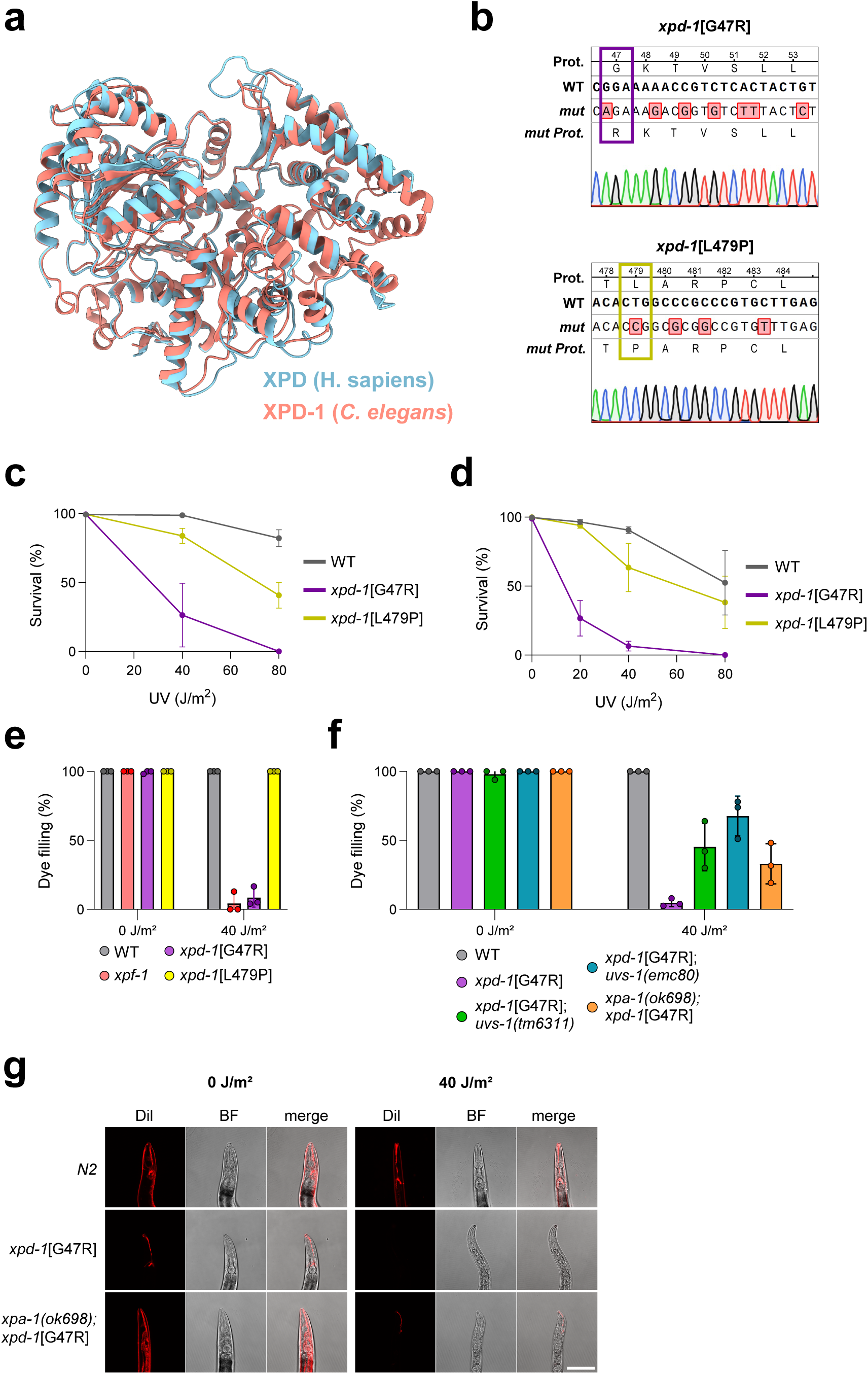
Futile DNA excision mediated by helicase-dead XPD causes severe neuronal dysfunction *in vivo in C. elegans*. **A** Structural superposition of human XPD (PDB: 6RO4, chain B; salmon) and *C. elegans* XPD-1 (AlphaFold model AF-Q9N3L2-F1, v6; sky blue). Structures were superposed in ChimeraX v1.6 using Matchmaker, yielding an RMSD of 1.059 ^°^A over 573 pruned C*α* atom pairs (1.765 ^°^A across all 669 pairs). **B** Sanger sequencing chromatograms of the *xpd-1* locus in *xpd-1* [G47R] and *xpd-1* [L479P] *C. elegans* strains, with the introduced codon change highlighted. **C** Germline and embryo survival of wild type, *xpd-1* [G47R] and *xpd-1* [L479P] *C. elegans* strains after UV-B irradiation (0, 40 and 80 J/m^2^). **D** L1 larvae survival of strains as in (**C**) after UV-B irradiation (0, 20, 40 and 80 J/m^2^). **E** Percentage of animals showing DiI dye-filling of chemosensory neurons for wild type (WT), *xpf-1*, *xpd-1* [G47R] and *xpd-1* [L479P] strains, scored 72 h after mock-treatment (0 J/m^2^) or 40 J/m^2^ UV-B irradiation. **F** Percentage of dye-filling animals as in (**E**) for wild type (WT), *xpf-1*, *xpd-1* [G47R], *xpd-1* [G47R]; *uvs-1(tm6311)*, *xpd-1* [G47R]; *uvs-1(emc80)* and *xpa-1(ok698)*; *xpd-1* [G47R] strains, mock-treated (0 J/m^2^) or irradiated with 40 J/m^2^ UV-B. **G** Representative images of DiI dye filling of chemosensory neurons for wild type (WT), *xpd-1* [G47R] and *xpa-1(ok698)*; *xpd-1* [G47R] animals from the experiment quantified in (**F**), scored 72 h after mock-treatment (0 J/m^2^) or 40 J/m^2^ UV-B irradiation. Scale bar, 100 *µ*m. Survival and dye-filling data (**C–F**) are mean and s.e.m. of three independent experiments. Source data are provided as a Source Data file.

We first assessed the functional impact of both G47R and L479P mutations on NER capacity in *C. elegans*, by testing the UV sensitivity of germ cells and embryos, which are largely dependent on GG-NER^78^. This showed that *C. elegans* G47R mutants are profoundly hypersensitive to UV irradiation, whereas L479P mutants are only mildly hypersensitive compared to wild type animals (Figure 5c). We next tested the UV sensitivity of L1 larvae, by quantifying UV-induced developmental arrest, which specifically depends on TC-NER capacity and the ability to process Pol II during TC-NER in neurons, as we had previously shown^36,78,79^.

This again showed that G47R mutants, but not L479P mutants, are extremely hypersensitive to UV irradiation (Figure 5d). These *in vivo* data faithfully reproduce our findings in human cells, confirming the differential severity of the mutations at the whole-organism level and showing that in particular the G47R mutation leads to severely compromised GG-NER and developmental arrest due to impaired neuronal TC-NER.

We then assessed and compared the impact of both XPD-1 mutations on neuron functionality after DNA damage induction, as progressive neurological decline is a major hallmark of CS pathology^28^. Functionally intact chemosensory neurons in the head of *C. elegans* are able to take up the fluorescent dye DiI from the environment through their ciliated dendrites^80^. We have previously shown that this ‘dye filling’ capacity reflects the functional integrity of these neurons after DNA damage induction. Dye filling depends on intact transcription maintained by TC-NER, and is impaired upon transcription inhibition or persistent TFIIH stalling such as occurs in UV-irradiated *xpf-1* mutants^21,79^. Therefore, we tested dye-filling capacity in UV-irradiated wild type, *xpf-1* (as control) and *xpd-1* mutant *C. elegans*. This showed that both wild type and L479P mutant neurons retained the ability to take up the fluorescent dye after UV irradiation (Figure 5e). In stark contrast, and similar to control *xpf-1* loss-of-function mutants, chemosensory neurons in almost all G47R mutant animals completely failed to take up the dye after UV irradiation, indicative of a severe, DNA damage-dependent functional defect in these neurons.

We tested if this functional defect was also observed in animals heterozygous for the G47R mutation, but found that neurons in these animals retained their capacity for dye filling after UV irradiation (Figure S4b). This indicates that helicase-dead XPD does not act in a dominant fashion, likely because in time the wild type XPD protein still present will facilitate the removal of all lesions. Subsequently, we investigated whether this severe UV-induced neuronal defect is due solely to the absence of DNA repair or whether, additionally, the futile DNA repair activity of G47R mutant XPD-1 contributes to this phenotype. Therefore, we tested whether the phenotype is alleviated by preventing the recruitment of mutant TFIIH. *C. elegans* neurons fully depend on TC-NER to maintain their transcriptional and cellular integrity upon UV-induced DNA damage induction^79^, which implies that any defects due to G47R mutant XPD-1 activity should depend on the TC-NER protein UVSSA, which is responsible for TFIIH recruitment^11,12^ and on the general NER protein XPA, which stimulates TFIIH DNA binding and activity^18,21^. We therefore crossed the G47R *xpd-1* mutant strain with previously generated TC-NER-deficient strains that lack the complete coding sequence of the *C. elegans* UVSSA ortholog *uvs-1* (referred to as *emc80* ), or lack the TFIIH-interacting motif in *uvs-1* (referred to as *tm6311* )^36,81^ and with a GG- and TC-NER-deficient strain that functionally lacks the *C. elegans* XPA ortholog XPA-1 (referred to as *ok698* )^76,82^. Remarkably, we observed that both *uvs-1* and *xpa-1* deficiency led to a substantial rescue of the dye-filling deficiency of G47R *xpd-1* mutant animals (Figure 5f–g), indicating that preventing the recruitment of mutant TFIIH alleviates this severe neuronal phenotype. These results provide strong evidence that *in vivo* the persistent futile DNA repair attempts initiated by helicase-deficient XPD are more detrimental to the functional integrity of cells than the mere absence of DNA repair.

## 4 Discussion

In this study, we compared the molecular NER mechanism of cells carrying different patient-derived XPD mutations associated with XP, XP/TTD or XPCS, to resolve the pathogenic mechanism underlying CS features. We show that G47R, an XPD helicase-dead mutation associated with the most severe clinical outcome of CS, elicits the most profound DNA damage sensitivity in human cells and *C. elegans*, while still supporting substantial NER-dependent DNA synthesis. Using an *in vivo* excision assay, we demonstrate that the NER machinery in G47R mutant cells excises normal sized oligonucleotides that are devoid of DNA damage. This excision without damage removal traps cells in a continuous futile repair cycle, as evidenced by the sustained UDS and failure to remove UV-induced photoproducts over time after UV irradiation. In *C. elegans*, this causes severe neuronal dysfunction after DNA damage induction, phenocopying the neuronal defects previously observed upon persistent TFIIH stalling in *xpf-1* and *xpg-1* null mutant animals^21^. Crucially, this neuronal dysfunction is largely rescued by preventing the TC-NER-dependent recruitment of mutant TFIIH. This demonstrates that the continuous futile repair cycle is more toxic than the mere absence of DNA repair in differentiated post-mitotic cells. Moreover, these findings provide clear *in vivo* mechanistic insight into the role of the XPD helicase activity during NER.

The central mechanistic question raised by these findings is how mutant XPD can support the assembly and activity of the NER incision machinery while failing to remove the DNA lesion that triggers the reaction. During NER, TFIIH is recruited after initial damage recognition and opens DNA by the translocase/helicase activities of XPB and XPD^83,84^. XPD translocates along the displaced single-stranded DNA in the 5*^′^ →* 3*^′^* direction until its progression is blocked by the lesion^16–18,85^. This physical blockage constitutes the damage verification step that precedes the endonucleolytic incisions made by ERCC1-XPF and XPG after opening of the DNA^22,23^. The G47R substitution sits within the conserved ATP-binding domain and abolishes XPD translocase activity^49,50^. As a result, TFIIH with helicase-dead XPD can be recruited to UV damage, as confirmed by our laser accumulation experiments (and observed before with other helicase-dead XPD mutants^85,86^), but cannot scan along the DNA to physically encounter the lesion. This decouples DNA damage verification from DNA incision, as the helicase-dead XPD is effectively locked in its arrested state from the moment of recruitment, licensing incision independently of where the lesion is positioned relative to TFIIH. This idea is supported by *in vitro* experiments showing that inhibiting TFIIH translocation, by ATP depletion or using a non-hydrolyzable ATP analog, permits XPG to incise a damage-free DNA substrate^22^. Consequently, ERCC1-XPF and XPG excise a fragment from the open bubble that does not contain the lesion (Figure 6). Subsequently, TFIIH is rapidly released and the resulting gap is filled in by DNA synthesis, while the lesion remains intact and triggers NER reinitiation as part of a continuous futile repair cycle. Consistent with this model, our FRAP data reveal that TFIIH with helicase-dead XPD is more mobile than wild type TFIIH after UV irradiation, indicating that it rapidly dissociates and does not form a stable, durable incision complex that remains stalled at the lesion^85^. Our observation that G47R mutant cells are resistant to trabectedin provides independent support for this model, as trabectedin normally blocks XPG during TC-NER at the incision step when it encounters the trabectedin-DNA adduct^72,73^. As a result, the single incision made by ERCC1-XPF is not repaired and this persistent DNA break becomes specifically toxic to TC-NER-proficient cells. Resistance of G47R mutant cells to trabectedin indicates that XPG does not encounter and is not blocked by the adduct, because DNA is incised at an incorrect position, i.e. adjacent to instead of flanking the lesion, and that the XPF-mediated break is therefore resolved. Together, these observations indicate that the XPD helicase activity is not strictly required for incision complex assembly and DNA incision, but is indispensable for the formation of a stable incision complex encompassing the lesion, such that DNA is correctly incised on both sides of the lesion.

**Figure 6.**
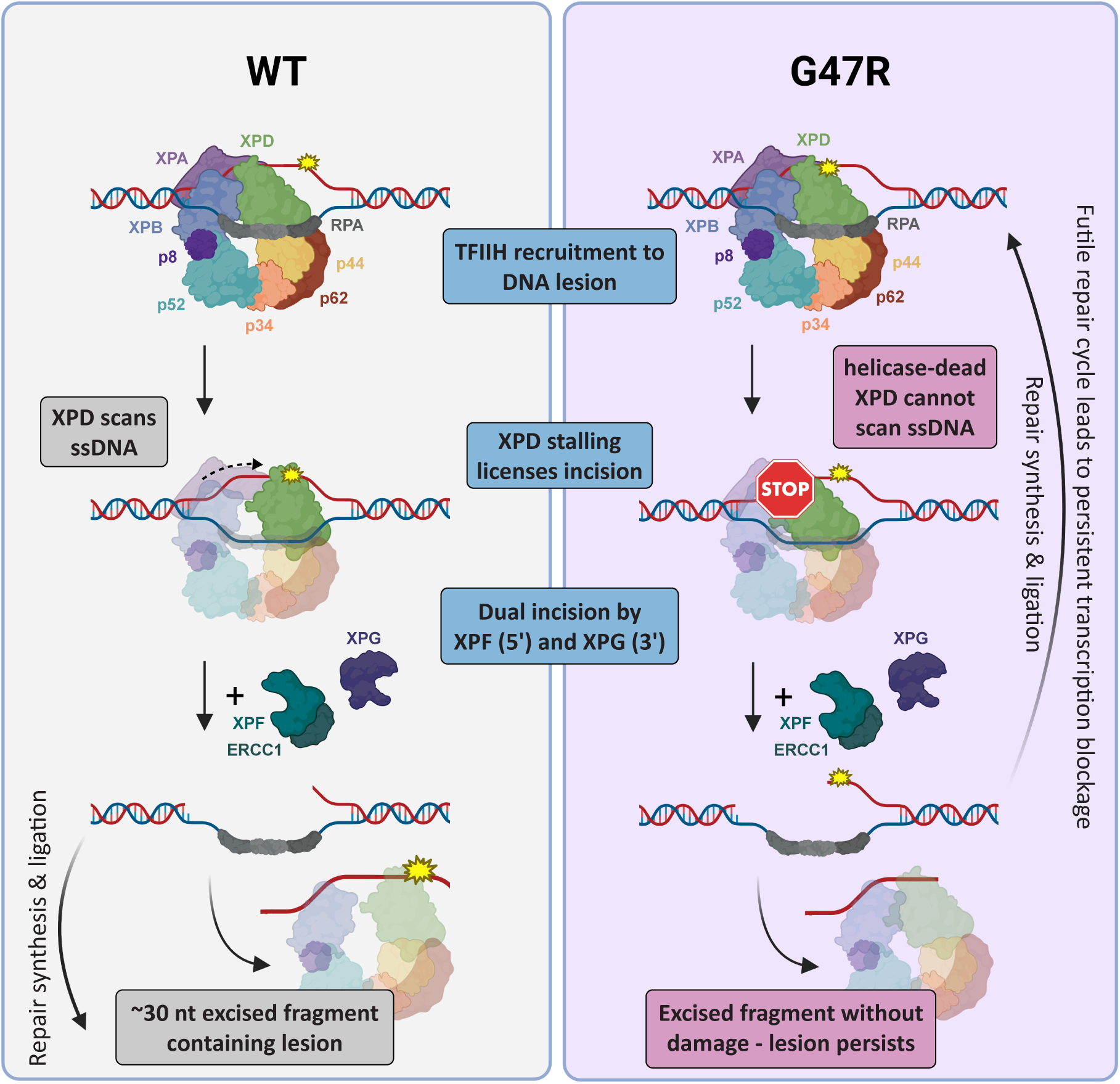
TFIIH encompassing helicase-dead XPD-G47R drives a futile NER cycle that causes persistent transcription blockage. Schematic comparison of nucleotide excision repair (NER) in wild type cells (left) and in XPD-G47R mutant cells (right). In wild type cells, TFIIH is recruited to the DNA lesion, where XPD scans the ssDNA and stalls at DNA damage to license dual incision by ERCC1-XPF (5*^′^*) and XPG (3*^′^*), producing an approximately 30 nt lesion-containing excised fragment, followed by gap-filling DNA repair synthesis and ligation. In G47R mutant cells, TFIIH is also recruited but the helicase-dead XPD cannot scan ssDNA, and, being stalled, licenses incision and excision of DNA to produce a damage-free fragment, leaving the original lesion in the genome. Continuous reinitiation of this unproductive cycle drives persistent transcription blockage. Created with BioRender.com.

Besides its role in DNA damage verification and incision, our results show that XPD heli-case activity promotes the clearance of lesion-stalled Pol II, in line with the idea that this facilitates backtracking and removal of Pol II from DNA damage to provide access for the downstream repair machinery^87,88^. We observed a partial defect in Pol II clearance from local DNA damage sites in G47R mutant cells, which is in agreement with a recent study that similarly showed that cells expressing the K48R or G675R helicase-dead XPD mutants display a partial Pol II clearance defect^37,59^. This study furthermore demonstrated that this defect is partial because Pol II can be cleared by a back-up extraction mechanism as well, involving the ubiquitin-dependent segregase VCP. CSB-mClover FRAP experiments confirmed that persistent stalling of Pol II at DNA lesions is not a dominant feature of G47R mutant cells. Importantly, the partial defect observed in G47R mutant cells is substantially less severe than the complete clearance defect observed in CSB knockout cells, showing that CSB itself is required for complete removal of lesion-stalled Pol II, as also noted before^35,59^. The partial Pol II clearance defect may therefore be a contributing factor to the mutant XPD phenotype, but it is the continuous futile DNA repair cycle that distinguishes XPD helicase mutant cells from cells with other XPD mutations, or with mutations in other GG-NER or TC-NER factors.

Our findings thus reveal a new and distinct mechanism by which persistent NER intermediates can drive CS pathology. Prior work has established that the persistent stalling of Pol II at DNA lesions underlies the severe CS symptoms in patients with mutations in TC-NER genes *CSB* and *CSA*^31,35–37,89,90^. This explains the difference in severity of symptoms compared to the clinically much milder UV^S^S, primarily caused by mutations in the TC-NER gene *UVSSA*. In *CSB* and *CSA* deficient cells, Pol II is not removed from DNA damage and therefore shields any means of repair, whereas in UV^S^S cells, Pol II is still efficiently cleared and repair can take place via alternative pathways so that transcription can recover. Moreover, CSB and CSA, but not UVSSA, were implicated in the transcription-coupled repair of DNA-protein crosslinks, implying that in *CSB* and *CSA* deficient cells these relatively large transcription blockages persist as well^38–40^. We furthermore showed that the more severe CS symptoms observed in certain *XPF* or *XPG* deficient patients can be attributed to a persistent repair intermediate as well, by showing that the persistent stalling of TFIIH in XPF or XPG-deficient cells inhibits transcription recovery and functionally impairs human and *C. elegans* cells^21,41^. Here, we identify a mechanistically distinct situation in which TFIIH with helicase-dead XPD does not persistently stall, but continues to engage in a repetitive futile manner with DNA damage, generating a different type of persistent transcription block. Each cycle of excision and DNA synthesis will transiently disrupt the DNA template and prevent transcription resumption at the damaged site. Recently, it was shown that cisplatin-induced neurotoxicity can be caused by a shortage of deoxynucleoside triphosphates (dNTPs) in neurons, which restricts DNA repair synthesis^91^. As a result, excessive but incomplete NER in response to cisplatin-treatment leaves behind persistent DNA gaps and breaks that trigger cell death. A similar problem may occur in XPD helicase-deficient cells with low dNTP pools, in which repetitive futile NER could lead to DNA breaks. Although mechanistically different, in each situation, the net outcome is prolonged transcriptional impairment, which explains why such molecularly different defects all lead to the same severe CS phenotype. Importantly, this shows that the persistent or recurrent NER intermediate is more toxic than the DNA lesion or absence of DNA repair itself, a principle that is most directly demonstrated by the partially restored neuronal function when the recruitment of G47R mutant TFIIH is prevented in *C. elegans*.

By specifically preventing TFIIH recruitment, through mutation of UVS-1 or XPA-1, we were able to substantially restore neuronal function in UV-irradiated G47R mutant animals. These results, together with our similar previous observations that preventing persistent Pol II or TFIIH stalling also significantly restores cellular function^21,36,41^, indicate that preventing the continuous engagement of the DNA repair machinery with DNA lesions could be a viable strategy for alleviating the very severe CS symptoms associated with certain NER deficiencies. It remains to be established whether such a strategy could be applied in mammalian model systems, and whether the selective modulation of TC-NER engagement could be therapeutically relevant to CS patients.

## 5 Materials and Methods

### Cell culture and generation

U2OS cells and patient-derived fibroblasts were cultured in DMEM (Lonza) supplemented with 1 % penicillin-streptomycin and 10 % or 15 % fetal calf serum (FCS; Serana), respectively. hTERT-RPE1 cells stably expressing inducible Cas9 (iCas9), generated previously^69^, were cultured in DMEM/F-12 (Lonza) supplemented with 1 % penicillin-streptomycin and 10 % FCS. All cells were maintained at 37 °C in a humidified 5 % CO_2_ atmosphere and routinely confirmed negative for mycoplasma contamination. All cell lines used in this study, including their origin and whether they were generated here or in previous publications, are listed in Supplementary Table 1. All chemicals and reagents used in this study, including catalogue numbers and suppliers, are listed in Supplementary Table 8.

Knock-in U2OS cell lines carrying XPD-G47R, XPD-L485P or XPD-S541R mutations were generated by nucleofecting Cas9 RNPs (Cas9 protein, tracrRNA, and crRNA) and ssODN repair templates (IDT) using the 4D-Nucleofector™ System (Lonza). Cells were transfected using the SE Cell Line 4D-Nucleofector™ XL Kit and program DC-100. To promote homology-directed repair, cells were cultured for 48 h post-transfection in medium supplemented with 2 *µ*M NU7441 (Selleckchem) and 10 *µ*M ART558 (MedChemExpress). Clonal lines were subsequently expanded and validated by PCR and Sanger sequencing. All sgRNA and ssODN sequences are listed in Supplementary Table 5.

For complementation experiments, hTERT-RPE1 iCas9 XPC^KO^ cells were transfected with wild type or mutant XPD-GFP variants, and endogenous XPD was subsequently depleted by crRNA-mediated CRISPR knock-down as described previously^59^. Briefly, cells were treated with 200 ng/ml doxycycline for 24 h to induce Cas9 expression, then transfected with 10 nM tracrRNA and 10 nM crRNA targeting an endogenous XPD sequence not present in the exogenous XPD-GFP construct, using RNAiMAX Lipofectamine (Thermo Fisher Scientific, 13778150). Three days after transfection, cells were collected and reseeded for the indicated experiments. Knock-down of endogenous XPD was verified by immunoblotting with an XPD antibody (ab54676, Abcam). crRNA sequences are listed in Supplementary Table 5.

For siRNA-mediated knock-down, cells were transfected with siRNA 48 h before each experiment using RNAiMAX (Invitrogen) according to the manufacturer’s instructions. Knock-down efficiency was confirmed by immunoblotting. All siRNA sequences are listed in Supplementary Table 4.

### Structural conservation mapping

Multiple sequence alignment of XPD orthologs from *Homo sapiens* (UniProt: P18074), *Mus musculus* (UniProt: O08811), *Bos taurus* (UniProt: A6QLJ0), *Arabidopsis thaliana* (UniProt: Q8W4M7), *Caenorhabditis elegans* (UniProt: Q9N3L2), and *Saccharomyces cerevisiae* (Rad3; UniProt: P06839) was performed using SnapGene software. Conservation scores derived from the alignment were mapped onto the cryo-EM structure of human TFIIH bound to DNA (PDB: 6RO4)^18^ using UCSF ChimeraX v1.6, with residues color-coded according to their degree of evolutionary conservation. The positions of patient mutations G47R, L485P, and S541R were annotated on the resulting structure. For the structural superposition of human XPD (PDB: 6RO4, chain B) and *C. elegans* XPD-1 (AlphaFold model AF-Q9N3L2-F1), the two structures were superposed in UCSF ChimeraX v1.6 using the Matchmaker tool. Pairwise sequence alignment of human XPD (UniProt: P18074) and *C. elegans* XPD-1 (UniProt: Q9N3L2) was performed with Clustal Omega and rendered with ESPript 3.0, using secondary structure elements derived from PDB 6RO4.

### Clonogenic survival assays

To assay survival, U2OS cells were seeded in triplicate in 6-well plates at a density of 500 cells per well. The following day, cells were treated with the indicated DNA-damaging agents. For UV survival assays, the culture medium was removed and cells were washed with PBS prior to exposure to UV-C light (254 nm), after which fresh medium was added.

For chemical treatments, cells were exposed to formaldehyde (FA; Thermo Fisher), illudin S (Gentaur), trabectedin (MedChemExpress), or 5-aza-2*^′^*-deoxycytidine (5-Aza-dC; Sigma-Aldrich). Treatment with FA, illudin S, and trabectedin was performed for 1 h, followed by three washes with PBS and replacement with fresh medium. Treatment with 5-Aza-dC was continuous until fixation.

After 7–10 days, colonies were fixed and stained with a solution of 0.1 % Coomassie Brilliant Blue (Sigma-Aldrich), 50 % ethanol, and 7 % acetic acid. Colonies were counted using the GelCount imager (Oxford Optronix). Colony numbers were normalized to mock-treated conditions (set to 100 %), and data were plotted as mean *±* s.e.m. The number of biological and technical replicates for each experiment is written in the corresponding figure legend.

### Immunofluorescence

For immunofluorescence, cells were seeded on 24 mm glass coverslips and fixed for 15 min in PBS containing 2 % formaldehyde and 0.1 % Triton X-100. Cells were permeabilized with PBS containing 0.1 % Triton X-100 and blocked with 1.5 % bovine serum albumin (BSA) in PBS for 30 min at room temperature. For experiments involving detection of UV-induced DNA lesions, DNA was first denatured with 70 mM NaOH in PBS for 10 min prior to blocking.

Primary antibodies were diluted in blocking buffer and incubated for 2 h at room temperature. Primary antibodies and their dilutions are listed in Supplementary Table 2, and secondary antibodies in Supplementary Table 3. Following three washes with PBS containing 0.1 % Triton X-100, cells were incubated with Alexa Fluor™-conjugated secondary antibodies (Invitrogen) and DAPI for 1 h at room temperature. After repeated washing, coverslips were mounted using Aqua-Poly/Mount (Polysciences). Digital images were acquired using a Zeiss LSM700 confocal microscope equipped with a 40*×* Plan-apochromat 1.3 NA oil-immersion lens (Carl Zeiss). Nuclear fluorescence intensities were quantified using Fiji/ImageJ software, with background fluorescence subtracted and values normalized to control conditions as specified per experiment.

### Immunofluorescence of elongating Pol II

To assess Pol II clearance from sites of UV-induced DNA damage, U2OS cells grown on coverslips were locally irradiated through 5 *µ*m pore filters (Millipore) with 100 J/m^2^ UV-C and subsequently incubated with 100 *µ*M 5,6-dichloro-1-*β*-D-ribofuranosylbenzimidazole (DRB; Sigma-Aldrich, D1916) for 1 or 2 h as indicated, to block new transcription initiation. Cells were then washed with PBS and fixed in 3.7 % formaldehyde in PBS for 15 min, followed by permeabilization in 0.5 % Triton X-100 in PBS for 10 min, both at room temperature. Non-specific binding was blocked by sequential incubation in 100 mM glycine in PBS for 10 min and 0.5 % BSA with 0.05 % Tween-20 in PBS for 10 min at room temperature. Cells were immunostained with primary antibodies against Pol II-S2P and CPD for 2 h at room temperature, followed by Alexa Fluor-conjugated secondary antibodies together with 0.1 *µ*g/ml DAPI for 1 h at room temperature. Antibodies and dilutions are listed in Supplementary Tables 2 and 3. Coverslips were mounted using Polymount (Brunschwig) and imaged on a Zeiss LSM700 confocal microscope equipped with a 40*×* Plan-apochromat 1.3 NA oil-immersion lens (Carl Zeiss).

### 6-4PP removal

To measure removal of UV-induced DNA photoproducts over time, U2OS cells grown on glass coverslips were globally irradiated with 4 J/m^2^ UV-C and fixed at 0, 3, 6, and 24 h post-irradiation. Immunofluorescence staining for 6-4PP, including NaOH denaturation, was performed as described above (see Immunofluorescence), and antibodies are listed in Supplementary Table 2. Nuclear fluorescence intensities were quantified using Fiji/ImageJ and normalized to the signal measured immediately after irradiation (0 h).

### UV-C laser accumulation

Accumulation of GFP-XPB at UV-C laser-induced DNA damage was measured in U2OS cells using a Leica TCS SP8 confocal microscope (LAS X software, v.3.3.0.16799, Leica) coupled to a 4.5 mW pulsed (15 kHz) diode-pumped solid-state 266 nm laser (Rapp Opto Electronic, Hamburg, Germany). Cells were grown on quartz coverslips and imaged through an Ultrafluar quartz 100*×*, 1.35 NA glycerol-immersion lens (Carl Zeiss Micro Imaging) at 37 °C and 5 % CO_2_. Fluorescence signals were background-corrected and normalized to the average pre-damage fluorescence levels in the irradiated region.

### Fluorescence Recovery After Photobleaching

FRAP was performed in U2OS cells expressing CSB-mClover using a Leica TCS SP8, and in U2OS cells expressing GFP-XPB using a Leica TCS SP5 confocal microscope, each equipped with an HC PL APO CS2 63*×*, 1.40 NA oil-immersion lens (Leica Microsystems). Cells were maintained at 37 °C and 5 % CO_2_ throughout imaging. Where indicated, cells were irradiated with UV-C prior to imaging. A strip of 512 *×* 16 pixels across the nucleus was imaged at 400 Hz using a 488 nm laser, with 0.2 s frame intervals for GFP-XPB and 0.4 s frame intervals for CSB-mClover. Fluorescence was measured for 5 pre-bleach frames to establish a steady-state level, followed by photobleaching of the strip using 1 frame at 100 % laser power. Fluorescence recovery was then recorded for 40 s (GFP-XPB) or 45 s (CSB-mClover). Fluorescence signals were background-corrected and normalized to the average pre-bleach fluorescence intensity. Immobile fractions (*F*_imm_) were calculated using the fluorescence intensity immediately after bleaching (*I*_0_) and the average steady-state fluorescence after complete recovery from untreated (*I*_final,unt_) and UV-C-treated cells (*I*_final,UV_), as previously described^92^.

### Immunoblotting

For immunoblotting of total cell extracts, cells were washed twice with PBS and lysed in RIPA buffer (50 mM Tris pH 7.5, 150 mM NaCl, 0.1 % SDS, 0.5 % sodium deoxycholate, 1 % NP-40) supplemented with EDTA-free protease inhibitor cocktail (Roche). Lysates were centrifuged for 10 min at 4 °C and mixed with Laemmli sample buffer, followed by boiling for 5 min at 95 °C. Proteins were separated on 4–15 % Mini-Protean TGX precast gels (Bio-Rad) and transferred onto PVDF membranes (0.45 *µ*m, Merck Millipore) overnight at 4 °C. Membranes were blocked for 1 h in 5 % BSA in PBS-T (0.05 % Tween-20) at room temperature and incubated with primary antibodies overnight at 4 °C. A complete list of primary antibodies and dilutions is provided in Supplementary Table 2. Following washing with PBS-T, membranes were incubated with CF™ 680- and CF™ 770-conjugated secondary antibodies (Sigma-Aldrich) and proteins were visualized using an Odyssey CLx infrared imaging system (LI-COR Biosciences). Image analysis was performed using Image Studio Lite (v.5.2, LI-COR).

### Unscheduled DNA synthesis

To measure global unscheduled DNA synthesis (UDS), U2OS cells grown on coverslips were incubated with 1 *µ*M Palbociclib (Selleck Chemicals) for 24 h to limit S-phase cells. Cells were then irradiated with 8 J/m^2^ UV-C and incubated for 1 h in medium containing 20 *µ*M 5-ethynyl-2*^′^*-deoxyuridine (EdU; Invitrogen). Where indicated, transcription was inhibited with 100 *µ*M DRB (Sigma-Aldrich, D1916), added 2 h before UV irradiation and maintained in the medium until fixation. Cells were fixed in 3.6 % formaldehyde in PBS and permeabilized with 0.1 % Triton X-100 in PBS for 10 min. EdU incorporation was visualized by incubating cells for 60 min at room temperature with a click-iT reaction cocktail containing 60 *µ*M Atto 594 Azide (Atto-Tec), 50 mM Tris-HCl pH 7.6, 4 mM CuSO_4_*·*5H_2_O (Sigma-Aldrich), and 10 mM ascorbic acid (Sigma-Aldrich). After washing with 0.1 % Triton X-100 in PBS, nuclei were stained with DAPI and coverslips were mounted onto microscope slides using Aqua-Poly/Mount (Polysciences). Fluorescent images were acquired using a Zeiss LSM700 confocal microscope equipped with a 40*×* Plan-apochromat 1.3 NA oil-immersion lens (Carl Zeiss). EdU signal intensities within nuclei were quantified using Fiji/ImageJ software, background fluorescence was subtracted, and values were normalized to control conditions.

To specifically measure transcription-coupled NER activity in hTERT-RPE1 cells expressing wild type or mutant XPD-GFP (see Cell culture and generation), cells were plated in DMEM supplemented with 8–10 % FCS and then serum-starved in DMEM without FCS for 24 h to reduce replicating cells and deplete the available deoxy-uridine pool. Cells were locally UV-irradiated through 5 *µ*m pore filters (Millipore, TMTP04700) with 100 J/m^2^ and immediately pulse-labeled with 20 *µ*M 5-ethynyl-2*^′^*-deoxyuridine (EdU; VWR) and 1 *µ*M FUdR (Sigma-Aldrich, F0503) for 4 h. After medium-chase with DMEM containing 10 *µ*M thymidine (Sigma-Aldrich, T1895) for 15 min, cells were fixed with 3.7 % formaldehyde in PBS for 15 min at room temperature and washed three times in PBS. Cells were then permeabilized for 20 min in PBS with 0.5 % Triton X-100 and blocked with 3 % BSA (Thermo Fisher) in PBS. EdU was visualized by click-iT chemistry, labeling for 30 min with a mix of 6 *µ*M Atto azide-Alexa 647 (Atto-Tec), 4 mM CuSO_4_ (Sigma-Aldrich), and 10 mM ascorbic acid (Sigma-Aldrich) in 50 mM Tris buffer (pH 8). Cells were then post-fixed with 2 % formaldehyde for 10 min and blocked with 100 mM glycine. After extensive washing with PBS, DNA was denatured with 0.5 M NaOH for 5 min, blocked with 10 % BSA in PBS for 15 min, and equilibrated in 0.5 % BSA and 0.05 % Triton X-100 in PBS (WB buffer). Locally damaged regions were visualized by staining for 2 h with anti-CPD (Cosmo Bio, 1:1000 in WB buffer), followed by Alexa Fluor 555-conjugated secondary antibody (Thermo Fisher, 1:1000 in WB buffer) for 1 h, counterstained with 0.1 *µ*g/ml DAPI, washed extensively with PBS and mounted in Polymount (Brunschwig).

### Recovery of RNA synthesis

To measure recovery of RNA synthesis (RRS) after UV-induced DNA damage, hTERT-RPE1 cells expressing wild type or mutant XPD-GFP (see Cell culture and generation) were grown on coverslips, irradiated with 12 J/m^2^ UV-C, and allowed to recover for the indicated times at 37 °C. Cells were then pulse-labeled for 1 h with 400 *µ*M 5-ethynyl-uridine (EU; Jena Bioscience) and washed for 15 min with DMEM without supplements. Cells were fixed with 3.7 % formaldehyde in PBS for 15 min, permeabilized with 0.5 % Triton X-100 in PBS for 10 min at room temperature, and blocked in 1.5 % BSA (Thermo Fisher) in PBS. EU incorporation was visualized by click-iT chemistry, labeling cells for 1 h with a solution containing 60 *µ*M Atto azide-Alexa 594 (Lumiprobe), 4 mM CuSO_4_ (Sigma-Aldrich), 10 mM ascorbic acid (Sigma-Aldrich), and 0.1 *µ*g/ml DAPI in 50 mM Tris buffer (pH 8). Cells were washed extensively with PBS before mounting in Polymount (Brunschwig).

### *In vivo* excision assay

NER excision activity was measured in U2OS cells as previously described^71^. Briefly, cells were irradiated with the indicated dose of UV-C and lysed in cold Triton X-100 lysis buffer (20 mM Tris-HCl pH 7.5, 150 mM NaCl, 1 mM EDTA, 1 mM EGTA, and 1 % Triton X-100), and incubated for 15 min on ice. Following centrifugation at 20,000 *×g* for 1 h, supernatants were treated with RNase A (Thermo Fisher) for 20 min at 37 °C, followed by proteinase K (Carl Roth) at 55 °C for 30 min. DNA was purified by phenol/chloroform extraction and ethanol precipitation, labeled using terminal deoxynucleotidyl transferase (New England Biolabs) and biotin-11-dUTP (Jena Bioscience), resolved on 10 % TBE-urea gels, and transferred to a nylon membrane. Biotin-labeled oligonucleotides were detected using HRP-conjugated streptavidin and chemiluminescent substrate (Thermo Fisher), imaged with an ImageQuant LAS 4000 Mini system, and quantified using ImageQuant TL software (GE Healthcare).

### Immunoprecipitation of excised oligonucleotides

To specifically detect 6-4PP-containing excision products, purified DNA was subjected to immunoprecipitation as previously described^71^. Protein G and anti-rabbit Dynabeads (Invitrogen) were precoupled with rabbit anti-mouse IgG (ab46540, Abcam; 1:1000) and anti-6-4PP antibody (Cosmo Bio) in IP buffer (20 mM Tris-HCl pH 8.0, 2 mM EDTA, 150 mM NaCl, 1 % Triton X-100, 0.5 % sodium deoxycholate) for 3 h at 4 °C. Beads were then incubated with purified DNA overnight at 4 °C, washed sequentially with four wash buffers of increasing stringency, and eluted in 50 mM NaHCO_3_ with 1 % SDS and 20 *µ*g/ml glycogen at 65 °C for 15 min. Eluted oligonucleotides were purified by phenol/chloroform extraction and ethanol precipitation, 3*^′^*-end labeled with biotin-11-dUTP, resolved on 10 % TBE-urea gels, and detected using HRP-conjugated streptavidin and chemiluminescence as described above.

### *C. elegans* strains and culture

*C. elegans* strains were cultured according to standard methods on nematode growth medium (NGM) agar plates seeded with *Escherichia coli* OP50 at 20 °C. All strains used in this study are listed in Supplementary Table 7. *xpd-1* mutant strains were generated by CRISPR-Cas9-mediated genome editing through germline microinjection of sgRNA and recombinant Cas9 protein together with an ssODN repair template; sgRNA sequences and repair templates are provided in Supplementary Table 6. Mutations were verified by genotyping PCR and Sanger sequencing.

### *C. elegans* survival assays

UV sensitivity of germ cells, embryos, and L1 larvae was assessed as previously described^78^. For germ cell and embryo survival, synchronized young adults were transferred to empty agar plates and irradiated with the indicated dose of UV-B (Philips TL-12 tubes, 40 W). Animals were then transferred to NGM plates seeded with OP50 bacteria and allowed to recover for 24 h, after which groups of 3–5 adults were transferred to fresh 6 cm plates and allowed to lay eggs for 2–3 h. Six plates per dose were used in each experiment. After 24 h, the number of unhatched eggs was scored relative to the total number of eggs laid to calculate the survival percentage.

For L1 larvae survival, eggs were collected from adult *C. elegans* by hypochlorite treatment and plated onto five technical replicate NGM agar plates per condition, seeded with HT115 bacteria. After 16 h, synchronized L1 larvae were irradiated with the indicated dose of UV-B (Philips TL-12 tubes, 40 W) and allowed to recover for 48 h, after which survival was scored by determining whether animals had developed beyond the L2 stage.

### Dye filling assay

To assay dye filling capacity, synchronized adult *C. elegans* were irradiated with the indicated dose of UV-B (Philips TL-12 tubes, 40 W) and allowed to recover on NGM culture plates seeded with OP50 bacteria for 72 h. Animals were then washed and incubated for 30 min in 10 *µ*g/ml DiI (Molecular Probes), diluted in M9 buffer from a stock solution in dimethylformamide (DMF). Subsequently, animals were allowed to recover for 1 h on culture plates, after which dye filling was scored using an Olympus SZX12 stereo microscope equipped with a U-RFL-T mercury lamp. Dye filling of heterozygous G47R *xpd-1* mutants was tested by crossing G47R mutant males with *unc-45(m94)* mutants and scoring dye filling of the non-Unc progeny.

### Statistical analysis

All data are presented as mean values with s.e.m. error bars unless otherwise indicated. Statistical analyses were performed using GraphPad Prism 10 for Windows (GraphPad Software, La Jolla, CA, USA). For comparisons across multiple groups, a one-way ANOVA was applied, using Dunnett’s multiple-comparison test for comparisons against a single reference condition and Tukey’s multiple-comparison test for comparisons among all conditions. The number of biological replicates and sample sizes for each experiment are reported in the respective figure legends. No statistical method was used to predetermine sample size. No data were excluded from the analyses. Experiments were not randomized, and investigators were not blinded during experiments or outcome assessment.

## 6 Data Availability

All data supporting the findings of this study are available within the article and its Supplementary Information file. Source data underlying all graphs and quantifications, as well as uncropped images of all blots and gels, are provided as a Source Data file with this paper. This study made use of publicly available structural and sequence data. The cryo-EM structure of human TFIIH bound to DNA is available in the Protein Data Bank under accession code 6RO4. Protein sequences were retrieved from UniProt under accession codes P18074 (*H. sapiens* XPD), O08811 (*M. musculus* XPD), A6QLJ0 (*B. taurus* XPD), Q8W4M7 (*A. thaliana* UVH6), Q9N3L2 (*C. elegans* XPD-1) and P06839 (*S. cerevisiae* Rad3). The predicted structure of *C. elegans* XPD-1 is available from the AlphaFold Protein Structure Database under model identifier AF-Q9N3L2-F1.

Cell lines and *C. elegans* strains generated in this study are listed in Supplementary Tables 1 and 7 and are available from the corresponding author upon reasonable request. Any remaining information is available from the corresponding author upon reasonable request.

## Code availability

No custom code or software was generated in this study. All analyses were performed using the commercially and publicly available software packages described in the Methods.

## 7 Acknowledgements

We thank the Erasmus MC Optical Imaging Center for microscope support. We thank Alan R. Lehmann for XPCS2, XP8BR and XPCS1PV patient fibroblasts, which were provided by the Genome Damage and Stability Centre Research Tissue Bank (GDSC-RTB), University of Sussex. We thank Karen Bauer and Thorsten Hoppe for *C. elegans unc-45* mutants. The CSB knockout U2OS cell lines were generated by Sidal Gündüz and Carlota Davó-Martínez and the XPC U2OS knockout by Ülkem Kaynak. Some *C. elegans* strains were provided by the *Caenorhabditis* Genetics Center (funded by NIH Office of Research Infrastructure Programs P40 OD010440) and the National Bioresource Project for the nematode. This work was financially supported by the Netherlands Organization for Scientific Research (OCENW.M20.343 and OCENW.XL.23.120), a National Research Council of Science & Technology (NST) grant from the Korea government (MSIT) (No. CAP22041-100), and the Korea Research Institute of Standards and Science (KRISS-2026–KP2026-0001). MSL was supported by the European Research Council Consolidator Grant STOP-FIX-GO (grant agreement No 101043815).

## Author contributions

DH, AFT and VvB generated cell lines and performed cell biology and imaging experiments. GHK and YK performed *in vivo* excision assays. PvdM performed TC-UDS and RRS assays. QY and AR performed cell survival experiments. DH, CVW, and KLT conducted *C. elegans* experiments. MSL, JAM, WV, JHC and HL conceptualized ideas and supervised experiments. DH and HL wrote the manuscript. All authors reviewed the manuscript.

---

**Figure S1. Sequencing and survival of XPD mutant cells. A** Sanger sequencing chromatograms of the *XPD* locus in XPD-G47R, -L485P and -S541R GFP-XPB knock-in U2OS cells, with the introduced codon change highlighted. **B** As in (**A**), for XPD-G47R, -L485P and -S541R CSB-mClover knock-in U2OS cells. **C** Representative confocal images of GFP-XPB fluorescence and DAPI in parental (WT) and XPD-G47R, -L485P and -S541R GFP-XPB knock-in U2OS cells. Scale bar, 10 *µ*m. **D** Clonogenic survival of parental (WT) and XPD-G47R, -L485P and -S541R GFP-XPB knock-in U2OS cells after UV-C treatment. Mean and s.e.m. of three independent experiments, each performed in technical triplicate. Source data are provided as a Source Data file.

**Figure S2. FRAP analysis in XPD mutant cells. A** FRAP curves of GFP-XPB in untreated and UV-irradiated (5–30 min after 10 J/m^2^) parental (WT) and XPC knockout GFP-XPB knock-in U2OS cells. **B–D** FRAP curves of GFP-XPB in untreated and UV-irradiated (5–30 min after 10 J/m^2^) XPD-G47R (**B**), XPD-L485P (**C**) and XPD-S541R (**D**) GFP-XPB knock-in U2OS cells. Immobile fractions were calculated and shown in Figure 2a. **E** FRAP curves of CSB-mClover in untreated and UV-irradiated (5–30 min after 10 J/m^2^) parental (WT), XPD-G47R and CSA knockout CSB-mClover knock-in U2OS cells. Immobile fractions were calculated and shown in Figure 2e. FRAP curves are normalized to pre-bleach fluorescence intensity. Each curve represents 10 cells from a representative experiment, except untreated XPD-G47R (13 cells) and untreated CSA knockout (9 cells). Source data are provided as a Source Data file.

**Figure S3. UDS in XPD helicase mutant cell lines and patient fibroblasts. A** Unscheduled DNA synthesis (UDS) measured by EdU incorporation for 1 h after 8 J/m^2^ UV-C irradiation in parental (WT) and XPD-G47R cells, in both the CSB-mClover and GFP-XPB knock-in U2OS backgrounds. EdU intensities were normalized to WT. n = 76 (WT, CSB-mClover), 78 (G47R, CSB-mClover), 105 (WT, GFP-XPB) and 119 (G47R, GFP-XPB) cells from two independent experiments. **B** UDS as in (**A**) in healthy and XPCS patient-derived fibroblasts expressing helicase-dead XPD (C5RO wild type control, XPCS2[G602D], XPCS1PV[R666W] and XP8BR[G675R/R669Gfs*40]), normalized to the wild type (C5RO) signal. n = 160 (C5RO), 141 (XPCS2), 128 (XPCS1PV) and 173 (XP8BR) cells from three independent experiments. **C** Immunoblot showing XPD-GFP expression and endogenous XPD silencing in the XPC knockout hTERT-RPE1 cells used for the RRS and TC-NER-UDS assays in Figure 3f–g. Conditions were untreated cells and cells transfected with crXPD without complementation or stably complemented with WT, G47R or K48R XPD-GFP. HDAC1 was used as loading control. A representative experiment is shown. Mean and s.e.m. are shown. Numbers represent p-values (one-way ANOVA corrected for multiple comparisons). Source data are provided as a Source Data file.

**Figure S4. Evolutionary conservation of *C. elegans* XPD-1 and dye-filling of heterozygous *xpd-1* mutants. A** Pairwise alignment of human XPD (UniProt: P18074) and *C. elegans* XPD-1 (UniProt: Q9N3L2), generated with Clustal Omega and rendered with ES-Pript 3.0. Identical residues are shown white-on-red; conservatively substituted residues are shown red-on-white, grouped using a BLOSUM62 matrix with an ESPript global score cutoff of 0.7. Secondary structure elements above the alignment are derived from the human structure (PDB: 6RO4). Human XPD and *C. elegans* XPD-1 share 448 identical residues across 761 aligned positions (58.9% identity). **B** Percentage of animals showing DiI dye filling of chemosensory neurons for *xpd-1* [G47R], *unc-45(m94)* and *unc-45(m94)*/+; *xpd-1* [G47R]/+ animals, scored 72 h after mock-treatment (0 J/m^2^) or 40 J/m^2^ UV-B irradiation. *unc-45* mutants were used to generate readily recognizable heterozygous animals, as described in the Methods. Source data are provided as a Source Data file.

