## Supplementary information for "Helicase-deficient TFIIH causes severe disease features via persistent DNA excision without damage removal"

Figure S1

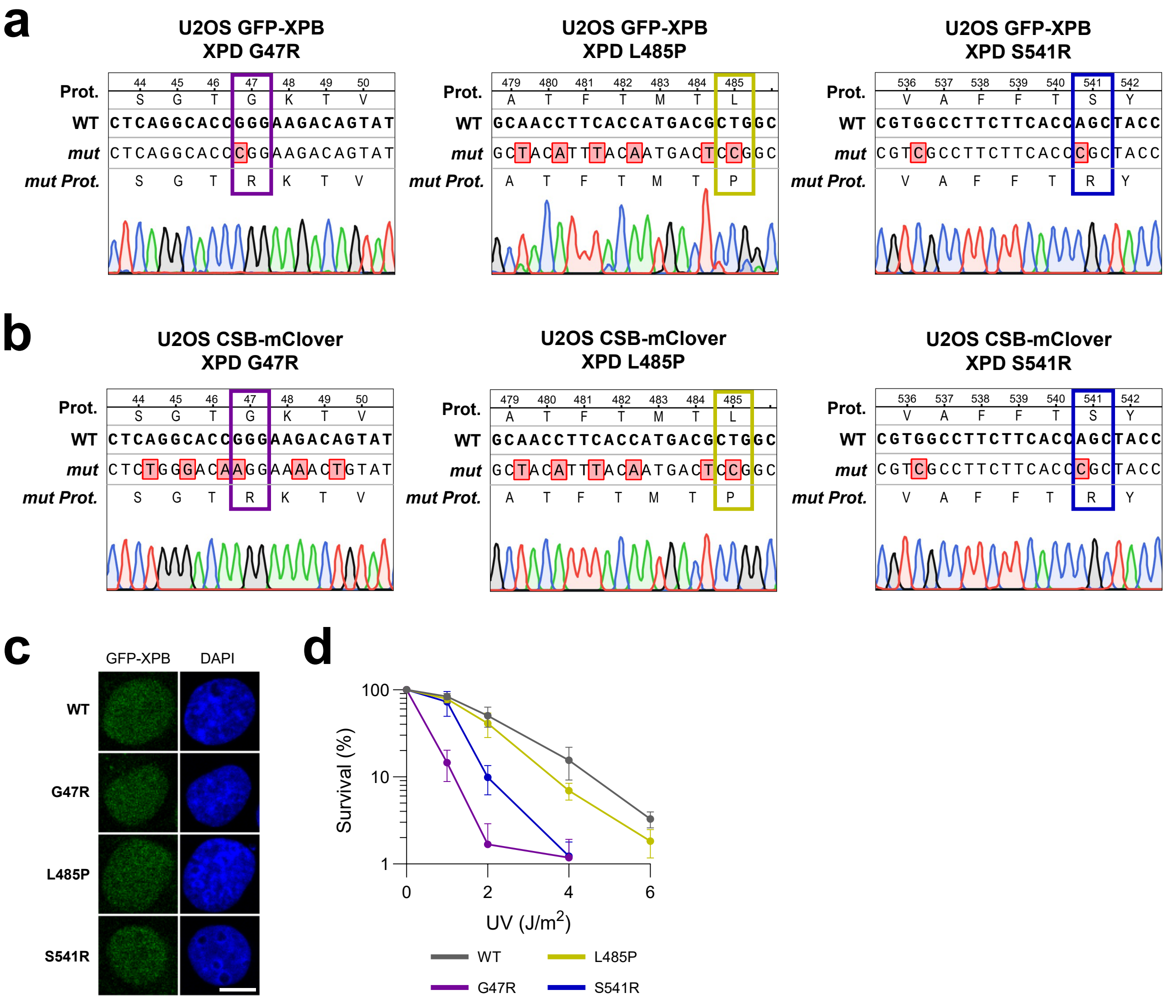
a
U2OS GFP-XPB
XPD G47R
U2OS GFP-XPB
XPD L485P
U2OS GFP-XPB
XPD S541R
Prot.
WT
mut
mut Prot.
b
U2OS CSB-mClover
XPD G47R
U2OS CSB-mClover
XPD L485P
U2OS CSB-mClover
XPD S541R
c
GFP-XPB
DAPI
WT
G47R
L485P
S541R
d
Survival (%)
UV (J/m2)
WT
L485P
G47R
S541R

Figure S2

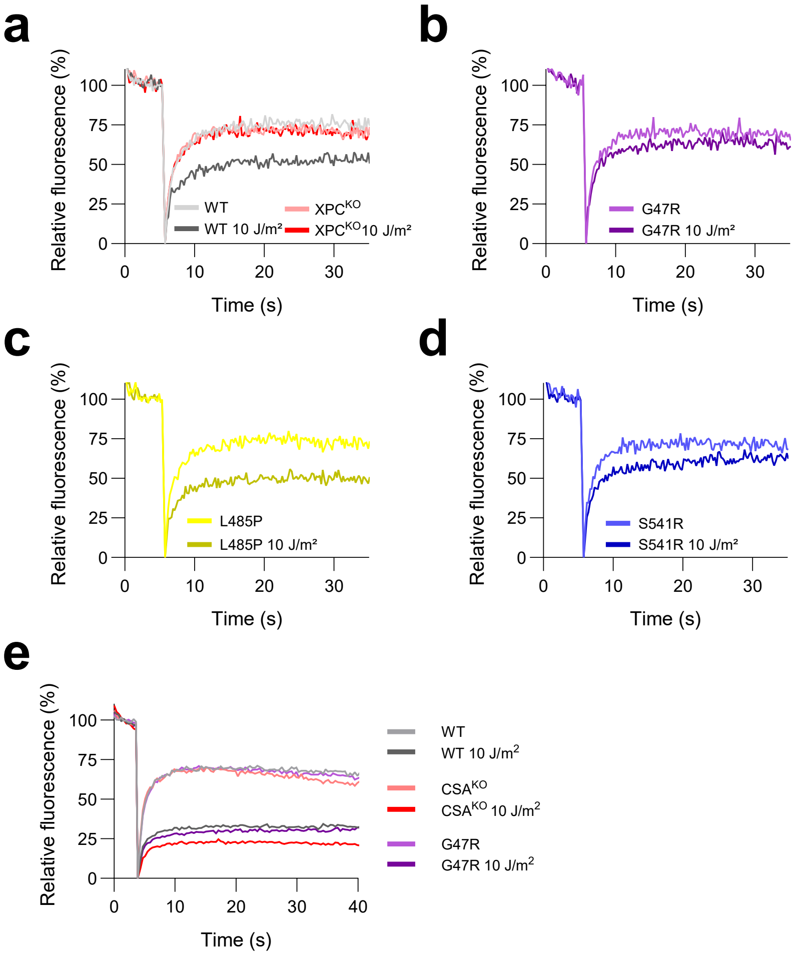
a
Relative fluorescence (%)
Time (s)
WT
WT 10 J/m²
XPC^KO
XPC^KO 10 J/m²
b
G47R
G47R 10 J/m²
c
L485P
L485P 10 J/m²
d
S541R
S541R 10 J/m²
e
WT
WT 10 J/m2
CSA^KO
CSA^KO 10 J/m2
G47R
G47R 10 J/m2

Figure S3

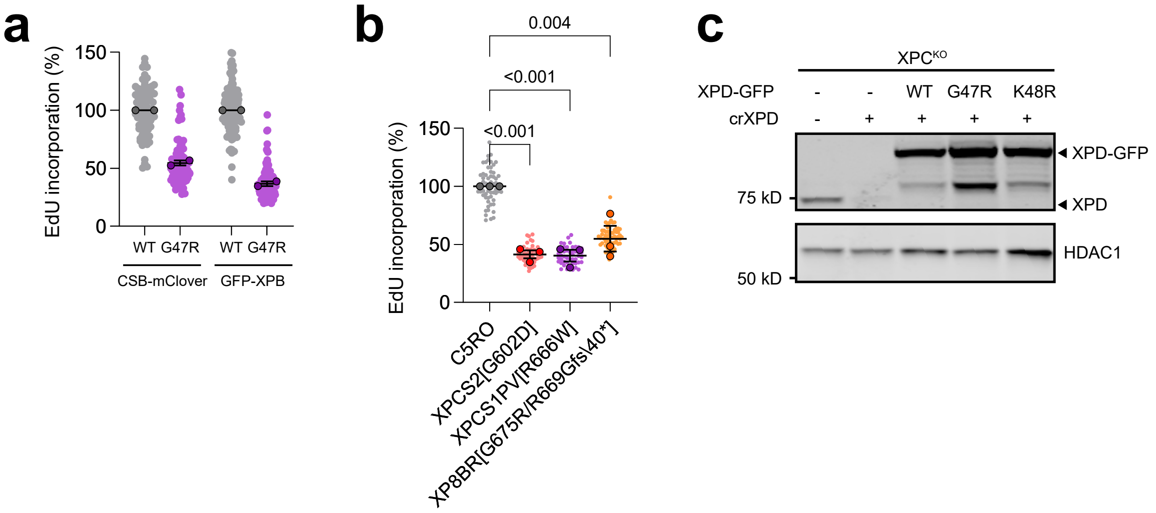

a
EdU incorporation (%)
150
100
50
0
WT G47R
WT G47R
CSB-mClover
GFP-XPB
b
0.004
<0.001
<0.001
EdU incorporation (%)
150
100
50
0
C5RO
XPCS2[G602D]
XPCS1PV[R666W]
XP8BR[G675R/R669Gfs\40*]
c
XPCKO
XPD-GFP - - WT G47R K48R
crXPD - + + + +
XPD-GFP
XPD
75 kD
HDAC1
50 kD

Figure S4

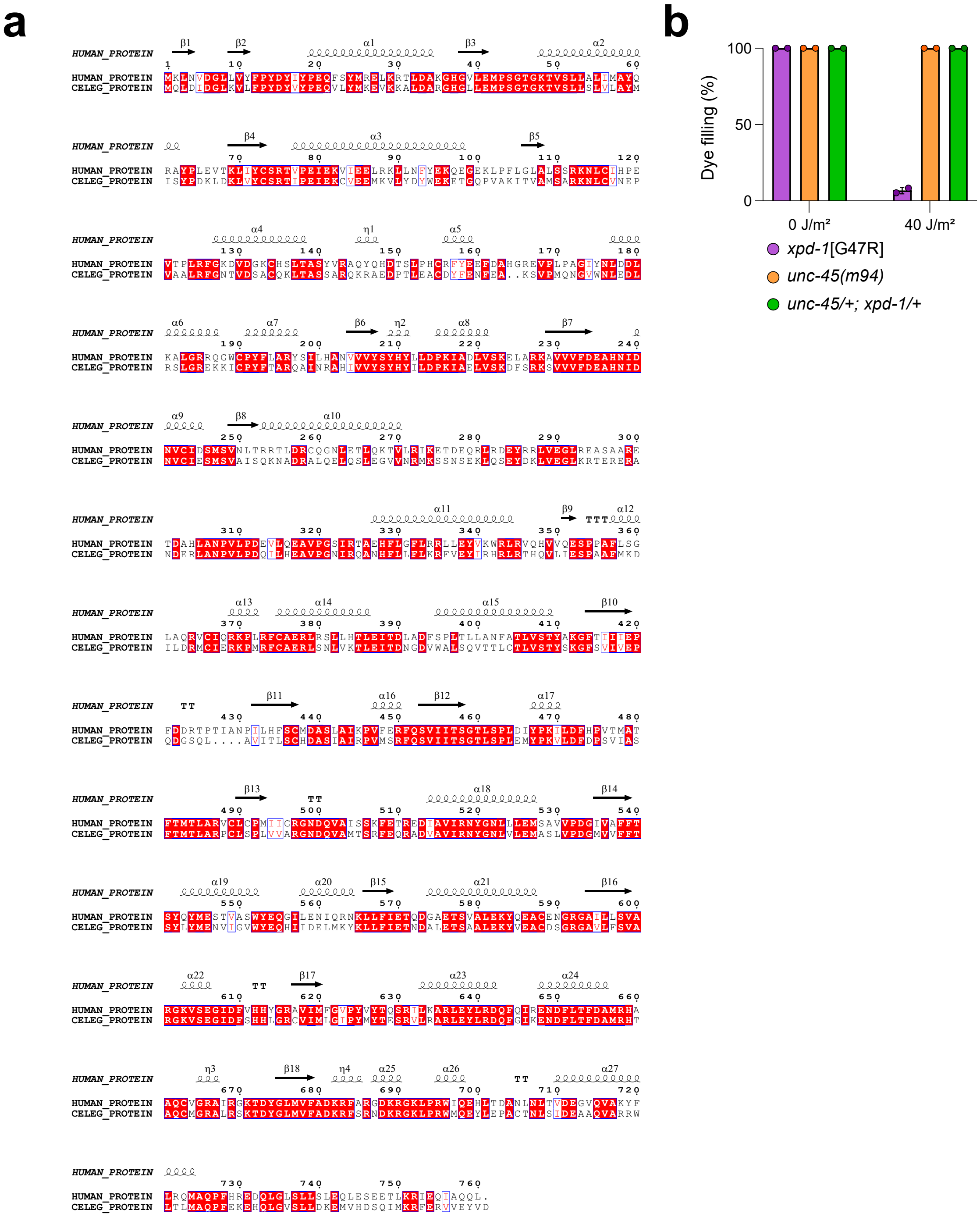
a
HUMAN_PROTEIN
CELEG_PROTEIN
b
Dye filling (%)
100
50
0
0 J/m²
40 J/m²
xpd-1[G47R]
unc-45(m94)
unc-45/+; xpd-1/+

### 9 Supplementary Tables

| Cell line | Source |
|---|---|
| U2OS | ATCC |
| U2OS GFP-XPB KI | [21] |
| U2OS CSB-mClover KI | [51] |
| U2OS GFP-XPB XPD-G47R KI | This study |
| U2OS GFP-XPB XPD-L485P KI | This study |
| U2OS GFP-XPB XPD-S541R KI | This study |
| U2OS CSB-mClover XPD-G47R KI | This study |
| U2OS CSB-mClover XPD-L485P KI | This study |
| U2OS CSB-mClover XPD-S541R KI | This study |
| U2OS CSB-mClover XPD-G47R KI XPA$^{KO}$ | This study |
| U2OS CSB-mClover CSA$^{KO}$ | [51] |
| U2OS XPC$^{KO}$ | This study |
| U2OS XPA$^{KO}$ | [21] |
| U2OS CSB$^{KO}$ | This study |
| hTERT-RPE1 iCas9 | [69] |
| hTERT-RPE1 iCas9 XPC$^{KO}$ | [59] |
| hTERT-RPE1 iCas9 XPC$^{KO}$ XPD-GFP$^{WT}$ | [59] |
| hTERT-RPE1 iCas9 XPC$^{KO}$ XPD-GFP$^{G47R}$ | This study |
| hTERT-RPE1 iCas9 XPC$^{KO}$ XPD-GFP$^{K48R}$ | This study |
| XPCS2 (XPCS-2BI) | [61] |
| XP8BR | [61] |
| XPCS1PV | [63] |
| C5RO | [93] |

Supplementary Table 1: Cell lines used in this study.

| Target | Cat. no. | Species | Supplier | Application | Dilution |
|---|---|---|---|---|---|
| XPB | ab190698 | Rabbit | Abcam | WB, IF | 1:1000 |
| XPA | GTX103168 | Rabbit | GeneTex | WB, IF | 1:500 |
| XPC | A301-112A | Rabbit | Bethyl | WB, IF | 1:1000 |
| XPD | ab54676 | Mouse | Abcam | WB | 1:1000 |
| XPF | sc-136153 | Mouse | Santa Cruz | WB, IF | 1:500 |
| XPG | A301-484A | Rabbit | Bethyl | IF | 1:1000 |
| RPA32 | GTX113004 | Rabbit | GeneTex | IF | 1:1000 |
| RNAPII-S2P | ab5095 | Rabbit | Abcam | IF | 1:1000 |
| 6-4PP | CAC-NM-DND-002 | Mouse | Cosmo Bio | IF, IP | 1:1000 |
| CPD | CAC-NM-DND-001 | Mouse | Cosmo Bio | IF | 1:1000 |
| HDAC1 | ab19845 | Rabbit | Abcam | WB | 1:1000 |

Supplementary Table 2: Primary antibodies used in this study.

| Specificity | Cat. no. | Conjugate | Supplier | Application |
|---|---|---|---|---|
| Goat anti-rabbit | A11034 | Alexa Fluor 488 | Invitrogen | IF |
| Goat anti-mouse | A11001 | Alexa Fluor 488 | Invitrogen | IF |
| Goat anti-mouse | A11029 | Alexa Fluor 488 | Thermo Fisher | IF |
| Goat anti-rabbit | A11012 | Alexa Fluor 594 | Invitrogen | IF |
| Goat anti-mouse | A11032 | Alexa Fluor 594 | Invitrogen | IF |
| Goat anti-rabbit | A21429 | Alexa Fluor 555 | Thermo Fisher | IF |
| Goat anti-mouse | A21422 | Alexa Fluor 555 | Thermo Fisher | IF |
| Goat anti-rabbit | SAB4600215 | CF™ 770 | Sigma-Aldrich | WB |
| Goat anti-rabbit | SAB4600200 | CF™ 680 | Sigma-Aldrich | WB |
| Goat anti-mouse | SAB4600214 | CF™ 770 | Sigma-Aldrich | WB |
| Goat anti-mouse | SAB4600199 | CF™ 680 | Sigma-Aldrich | WB |
| Rabbit anti-mouse | ab46540 | Unconjugated | Abcam | IP |

Supplementary Table 3: Secondary antibodies used in this study.

| Name | Sequence (5′–3′) | Catalogue no. |
|---|---|---|
| siCtrl | UGGUUUACAUGUUGUGUGA | D-001210-02-20 |
| siXPF | AAGACGAGCUCACGAGUAU | D-019946-04 |
| siExo1 | GCACGUAAUUCAAGUGAUG | Custom (Dharmacon) |

Supplementary Table 4: siRNA oligonucleotides used in this study.

| Target | Type | Sequence (5′–3′) |
|---|---|---|
| XPD G47R KI (CSB-mClover) | sgRNA | ACTGTCTTCCCGGTGCCTGA |
| XPD G47R KI (CSB-mClover) | ssODN | G*C*C*CCTCTGGTCCCCAACATGCAGG<br>GTCATGGAGTCCTGGAGATGCCCTCTG<br>GGACAAGGAAAACTGTATCCCTGTTG<br>GCCCTGATCATGGCATACCAGAGAGTG<br>AGTGATGC*G*C*T |
| XPD G47R KI (GFP-XPB) | sgRNA | CGGGAAGACAGTATCCCTGT |
| XPD G47R KI (GFP-XPB) | ssODN | T*G*T*GTGCCCAAGGTTCTGAGACCCT<br>GTGTGTTGCCCCTCTGGTCCCCAACAT<br>GCAGGGTCATGGAGTCCTGGAGATGC<br>CCTCAGGCACCCGGAAGACAGTATCCC<br>TGTTAGCCCTGATCATGGCATACCAGA<br>GAGTGAGTGATGCGCTGAACCCGTAA<br>AGGCAGACAAAGGAAGGGGCGGGACA<br>GGGACTGAGTCCG*C*T*T |
| XPD L485P KI | sgRNA | GGCAACCTTCACCATGACGC |
| XPD L485P KI | ssODN | C*C*G*CTGGACATCTACCCCAAGATCC<br>TGGACTTCCACCCCGTCACCATGGCTA<br>CATTTACAATGACTCCGGCACGGGTCT<br>GCCTCTGCCCTATGGTGAGTGGGAGA<br>GGCTAGGGCT*G*G*G |
| XPD S541R KI | sgRNA | TGTGGTCCCTGATGGCATCG |
| XPD S541R KI | ssODN | G*A*G*AGGGCCCAACCTCTGACCCCTT<br>GCAGCTGTGATCCGGAACTATGGGAA<br>CCTCCTGCTGGAGATGTCCGCTGTGGT<br>CCCTGATGGCATCGTCGCCTTCTTCAC<br>CCGCTACCAGTACATGGAGAGCACCGT<br>CGCCTCCTGGTATGAGCAGGTACGCCT<br>GGCCACCCCCTCCCTGCACCTGCTCTC<br>CTCAGTCCCTG*G*C*A |
| XPD knockdown (RPE1) | crRNA | AAGGAACAGGTGCTCACCTC |

Supplementary Table 5: sgRNA, crRNA, and ssODN sequences used for CRISPR-Cas9 genome editing and crRNA-mediated knockdown in human cell lines.

| Target | Type | Sequence (5′–3′) |
|---|---|---|
| *xpd-1* G47R KI | sgRNA | CACCAATGACAGTAGTGAGA |
| *xpd-1* G47R KI | ssODN | C*A*G*GGTCATGGACTACTGGAAATGCCGTCAGGA ACCAGAAAGACGGTGTCTTTACTCTCATTGGTGTT GGCTTATATGATATCGTATCCGGAT*A*A*G |
| *xpd-1* L479P KI | sgRNA | CTACGAGGGGACTCAAGCAC |
| *xpd-1* L479P KI | ssODN | C*G*A*CCCGTCCGTCATCGCATCATTCACAATGACA CCGGCGCGGCCGTGTTTGAGCCCTCTCGTCGTTGC ACGTGGAAATGACCAAGTGGCGATGAC*G*T*C |

Supplementary Table 6: sgRNA and ssODN sequences used for CRISPR-Cas9 genome editing in *C. elegans*.

| Strain ID | Genotype | Source |
|---|---|---|
| N2 | *C. elegans* Bristol N2 | CGC |
| HAL260 | *gtf-2H1(emc202[AID::GFP::gtf-2H1])* IV | [36] |
| HAL601 | *xpd-1(emc301[G47R])* III; *gtf-2H1(emc202[AID::GFP::gtf-2H1])* IV | This study |
| HAL604 | *xpd-1(emc304[L479P])* III; *gtf-2H1(emc202[AID::GFP::gtf-2H1])* IV | This study |
| HAL270 | *xpf-1(tm2842)* II | [94] |
| HAL257 | *uvs-1(emc80)* V | [36] |
| HAL121 | *uvs-1(tm6311)* V | [36] |
| HAL607 | *xpd-1(emc301[G47R])* III; *gtf-2H1(emc202[AID::GFP::gtf-2H1])* IV; *uvs-1(emc80)* V | This study |
| HAL605 | *xpd-1(emc301[G47R])* III; *gtf-2H1(emc202[AID::GFP::gtf-2H1])* IV; *uvs-1(tm6311)* V | This study |
| GJ1566 | *xpa-1(ok698)* I | [82] |
| HAL903 | *xpa-1(ok698)* I; *xpd-1(emc301[G47R])* III; *gtf-2H1(emc202[AID::GFP::gtf-2H1])* IV | This study |
| PP32 | *unc-45(m94)* III | K. Bauer & T. Hoppe |

Supplementary Table 7: *C. elegans* strains used in this study.

| Reagent | Cat. no. | Supplier |
|---|---|---|
| 5-ethynyl-2′-deoxyuridine (EdU) | A10044 | Invitrogen |
| 5-ethynyl-2′-deoxyuridine (EdU) | CLK-N001-100 | VWR |
| 5-ethynyl uridine (EU) | CLK-N002-10 | Jena Bioscience |
| 5-fluoro-2′-deoxyuridine (FUdR) | F0503 | Sigma-Aldrich |
| Thymidine | T1895 | Sigma-Aldrich |
| Atto 594 Azide | 5B930 | Lumiprobe |
| Atto 594 Azide | AD594-105 | Atto-Tec |
| Atto 647 Azide | AD647-105 | Atto-Tec |
| Palbociclib | S1116 | Selleck Chemicals |
| Doxycycline | D9891 | Sigma-Aldrich |
| 4′,6-diamidino-2-phenylindole (DAPI) | D9542 | Sigma-Aldrich |
| Aqua-Poly/Mount | 18606-20 | Polysciences |
| DRB | D1916 | Sigma-Aldrich |
| Formaldehyde | 28906 | Thermo Fisher |
| Illudin S | I019 | Gentaur |
| Trabectedin | HY-50936 | MedChemExpress |
| 5-aza-2′-deoxycytidine (5-Aza-dC) | A3656 | Sigma-Aldrich |
| Lipofectamine RNAiMAX | 13778500 | Invitrogen |
| Lipofectamine RNAiMAX | 13778150 | Thermo Fisher |
| NU7441 | S2638 | Selleckchem |
| ART558 | HY-141520 | MedChemExpress |
| Terminal deoxynucleotidyl transferase | M0315S | New England Biolabs |
| Biotin-11-dUTP | NU-803-BIOX-S | Jena Bioscience |
| Protein G Dynabeads | 10003D | Invitrogen |
| Anti-rabbit Dynabeads | 11203D | Invitrogen |
| RNase A | EN0531 | ThermoScientific |
| Proteinase K | 7528.4 | Carl Roth |
| 2× Laemmli sample buffer | S3401 | Sigma-Aldrich |
| Isopore membrane filters (5 $\mu$m, TMTP) | TMTP04700 | Millipore |
| DiI | D3911 | Molecular Probes |

Supplementary Table 8: Chemicals and reagents used in this study.
